# A One-Shot Multivalent Live-Attenuated Candidate Influenza Vaccine against Divergent Zoonotic H5N1 Clades

**DOI:** 10.64898/2026.08.02.742385

**Authors:** Ahmed M. Elsayed, Ramya S. Barre, Arash Rahmani, Ruby A. Escobedo, Trushar Jeevan, Himadri Nath, Esteban M. Castro, Vinay Shivanna, Aitor Nogales, James Kobie, Richard J. Webby, Elsayed M. Abdelwhab, Adolfo García-Sastre, Luis Martinez-Sobrido

## Abstract

The continued emergence of genetically diverse high pathogenicity avian influenza (HPAI) H5N1 viruses with zoonotic potential highlights the urgent need for developing vaccines capable of providing broad protection against multiple circulating clades. Here, we developed a one-shot, multivalent, live-attenuated influenza vaccine (LAIV) based on the temperature-sensitive (ts), cold-adapted (ca), and attenuated (*att*) influenza A/Ann Arbor/6/1960 master donor virus (MDV) that incorporates the hemagglutinin (HA) and neuraminidase (NA) glycoproteins from representative clades 2.3.4.4b (A/Louisiana/12/2024), 2.3.2.1a (A/Victoria/149/2024), and 2.3.2.1e (A/Cambodia/2302009/2023) H5N1 viruses. A single intranasal (IN) immunization of C57BL/6 mice with the multivalent LAIV elicited robust humoral immune responses, with immune sera exhibiting broad cross-reactivity against antigens from all three H5N1 clades included in the vaccine. Following homologous viral challenge, vaccinated C57BL/6 mice were completely protected from disease, demonstrating the immunogenicity and protective efficacy of the multivalent LAIV. By simultaneously targeting antigenically distinct H5N1 lineages with pandemic potential, this strategy expands antigenic coverage within a single LAIV to confirm pan-H5N1 protection. Together, these findings support the development and implementation of this multivalent LAIV as a broadly protective pan-H5N1 LAIV for pandemic preparedness.

**Significance:** The increasing genetic diversity of zoonotic H5N1 viruses complicates vaccine design. We developed a multivalent live-attenuated influenza vaccine (LAIV) based on the temperature-sensitive, cold-adapted, and attenuated (*ts*, *ca*, *att*) master donor virus (MDV) influenza A/Ann Arbor/6/1960 backbone that expresses the hemagglutinin (HA) and neuraminidase (NA) glycoproteins of H5N1 clades 2.3.4.4b, 2.3.2.1a, and 2.3.2.1e. A single intranasal (IN) immunization with the multivalent LAIV induced broadly cross-reactive neutralizing antibody (NAb) responses and protected experimental vaccinated animals against homologous lethal viral challenge, demonstrating the feasibility of the multivalent LAIV to protect against H5N1 clades of highest concern to humans. These findings demonstrate the feasibility of developing and implementing this multivalent LAIV as a broad protective pan-H5N1 LAIV against divergent H5N1 viruses for human use.

## Introduction

Influenza viruses are enveloped, segmented, single-stranded, negative-sense RNA viruses that belong to the family *Orthomyxoviridae*. Compared to influenza B, C, and D genera, influenza A viruses (IAVs) have a wider host range, which includes humans, other mammals, and birds. Each year, influenza virus infections account for approximately 3-5 million cases and around 290,000-650,000 fatalities in humans worldwide (1). IAVs remain a major public health threat, causing seasonal epidemics and occasional pandemics with substantial morbidity and mortality (2). The continued emergence of human IAVs is driven by two key evolutionary mechanisms. First, novel IAVs can arise through genetic reassortment, introducing antigenically distinct viruses into immunologically naïve human populations (3, 4). Second, once established in humans, IAVs undergo continuous antigenic evolution that facilitates viral fitness and sustains human-to-human transmission even in the presence of pre-existing immunity (2). However, despite sporadic human infections, these viruses are currently considered to pose a low human pandemic risk because of the lack of sustained human-to-human transmission (5).

Avian influenza viruses (AIVs) are contagious and primarily affect poultry and wild birds. AIVs are classified as high or low pathogenic viruses (HPAIV and LPAIV, respectively) depending on the molecular characteristics of the virus and their ability to cause systemic disease and mortality in chickens (2). HPAIV are associated with H5 and H7 subtypes of hemagglutinin (HA) surface glycoprotein. Disturbingly, HPAIV outbreaks in poultry are no longer an occasional phenomenon and the range of wild bird and mammal species affected by HPAIV are expanding, leading to HPAIV strains containing genetic markers of adaptation to mammal hosts (6–8). Animal-to-human transmission of HPAIV has occasionally occurred, while limited or rare transmissions among humans have been reported (9). Of these, HPAIV H5N1 stand out due to their higher case-fatality rates (CFR) in humans, posing concerns for public health. A recent example is the emergence of HPAIV H5N1 in dairy cattle in the United States of America (USA) in 2024, highlighting the ongoing continuous risk of zoonotic spillover (3, 4). In addition, H6, H7, H9, and H10 AIV subtypes have caused zoonotic infections in humans and infected other mammals (2). Since the first recognized human infection in 1997, HPAIV H5N1 has caused approximately 993 sporadic laboratory-confirmed human infections, including 477 fatalities, associated with multiple H5N1 clades across more than 20 countries resulting in an overall CFR of 48% (10). Peaks in human H5N1 cases occurred in 2006 (115 cases, 9 countries), 2015 (145 cases, 4 countries) and 2024 (81 cases, 5 countries) (10). Recent outbreaks of H5N1 infections associated with morbidity and mortality in humans and other mammals such as minks, cats, and seals have been reported in Cambodia, Australia, and USA (4, 6–8, 10–12). These infections were associated with H5N1 clades 2.3.4.4b (USA), 2.3.2.1a (Australia) and 2.3.2.1e (Cambodia) (13–16).

Vaccination remains the primary and most effective strategy to protect humans and animals against infections, including influenza (17–21). Three types of vaccines have been approved for use in humans to prevent seasonal influenza infections: inactivated influenza vaccines (IIV), recombinant HA vaccines, and live-attenuated influenza vaccines (LAIV) (19, 20). IIV have been shown to be ∼60–90% effective in reducing morbidity and mortality associated with seasonal influenza infections by inducing humoral immune response towards the surface viral HA glycoprotein and to a less and variable extent to the neuraminidase (NA) glycoprotein. However, IIV induce limited T cell-mediated immune responses (21, 22). In contrast, LAIV are intranasally (IN) administrated and mimic the natural route of IAV infection. They induce innate immunity and broad adaptive immune responses that resemble those induced during natural influenza viral infection, including mucosal cellular and humoral responses (22–26). Importantly, LAIV can prime specific T cell-mediated immune responses in naive populations and provide cross-protection against heterosubtypic strains. This LAIV protection profile is a highly desirable requisite for effective influenza vaccine candidates.

Although candidate vaccine viruses (CVVs) for the development of IIV to prevent AIV infections in humans, including H5N1, have been developed (27), they are not ready yet for widespread use. CVVs are based on the backbone of the high-growth influenza A/Puerto Rico/8/1934 H1N1 (PR8) harboring the HA and NA genes from the IAV strains recommended by the World Health Organization (WHO). Although LAIV could have high efficacy, there is no LAIV currently approved for use in humans or poultry, for the prevention of AIVs, including H5N1 infections. The genetic background of most licensed LAIV was developed by incorporating the internal segments (PB2, PB1, PA, NP, M, and NS) of the master donor virus (MDV) influenza A/Ann Arbor/6/1960 H2N2 (AA) with the surface HA and NA glycoproteins of circulating IAVs. LAIV presents with a cold-adapted (*ca*), temperature-sensitive (*ts*), and attenuated (*att*) phenotype of the MDV AA that replicates efficiently at low but not high temperatures (*ts* and *ca*) with a safety profile (*att*) (28, 29). Here, we developed a multivalent LAIV based on the *ts*, *ca*, and *att* AA backbone that targets HPAIV H5N1 clades 2.3.4.4b, 2.3.2.1a, and 2.3.2.1e. A single IN immunization with this multivalent LAIV elicited robust and durable protective immune responses against HPAIV H5N1 viruses and has the potential to provide pan-H5N1 protection against currently and newly emerging H5N1 variants with high zoonotic potential within these clades.

## Results

### Cross-reactivity of mAbs and ferret sera induced by CVV vaccination against clades 2.3.4.4b, 2.3.2.1a, and 2.3.2.1e H5 and N1 proteins

To assess the antigenic cross-reactivity of the three PR8-based H5N1 viruses expressing the HA and NA of A/Louisiana/12/2024 (clade 2.3.4.4b), A/Victoria/149/2024 (clade 2.3.2.1a), or A/Cambodia/2302009/2023 (clade 2.3.2.1e), IFA was performed using a panel of anti-H5 and anti-N1 monoclonal antibodies (mAbs) (**Fig. 1**). Infected cells were identified by staining with an anti-IAV NP pAb, while H5 HA or N1 NA expressions were detected using the indicated anti-H5 (**Fig. 1A**) and anti-N1 (**Fig. 1B**) mAbs. Interestingly, the three PR8-based H5N1 clade viruses (**Fig. 2A**) exhibited distinct Ab recognition profiles. The reference mAbs VN04-2, VN04-10, VN04-13, and VN04-16 showed stronger cross-reactivity with PR8H5N1c21a and PR8H5N1c21e, whereas little reactivity was observed with PR8H5N1c44b. In contrast, the anti-H5 mAb 464E11 recognized PR8H5N1c44b and PR8H5N1c21a but did not react, at least to the same extent, with PR8H5N1c21e. A similar pattern was observed for mAb 6B4, which detected PR8H5N1c44b and PR8H5N1c21a but failed to clearly recognize PR8H5N1c21e. By comparison, mAbs VN04-8, VN04-9 and 23E6 reacted similarly with all three viruses, suggesting that these mAbs target antigenic epitopes that are conserved across the tested H5 clade viruses. In contrast, mAbs 11F4 and 1C10 showed little or no detectable binding to any of the three viruses under the same experimental conditions (**Fig. 1A**). In parallel, distinct recognition patterns were also observed among the NA-specific mAbs. CD6 recognized PR8H5N1c44b and PR8H5N1c21e but did not react with PR8H5N1c21a. In contrast, 2B9 could slightly react with all three PR8-based H5N1 viruses with robust recognition of PR8H5N1c21e, while 109012 could strongly detect all three PR8-based H5N1 viruses with stronger recognition of PR8H5N1c44b and PR8H5N1c21a, suggesting that they target epitopes are, to some extent, conserved across these strains. By comparison, 3A2 failed to recognize any of the three viruses under the experimental conditions tested (**Fig 1B**).

**Figure 1.**
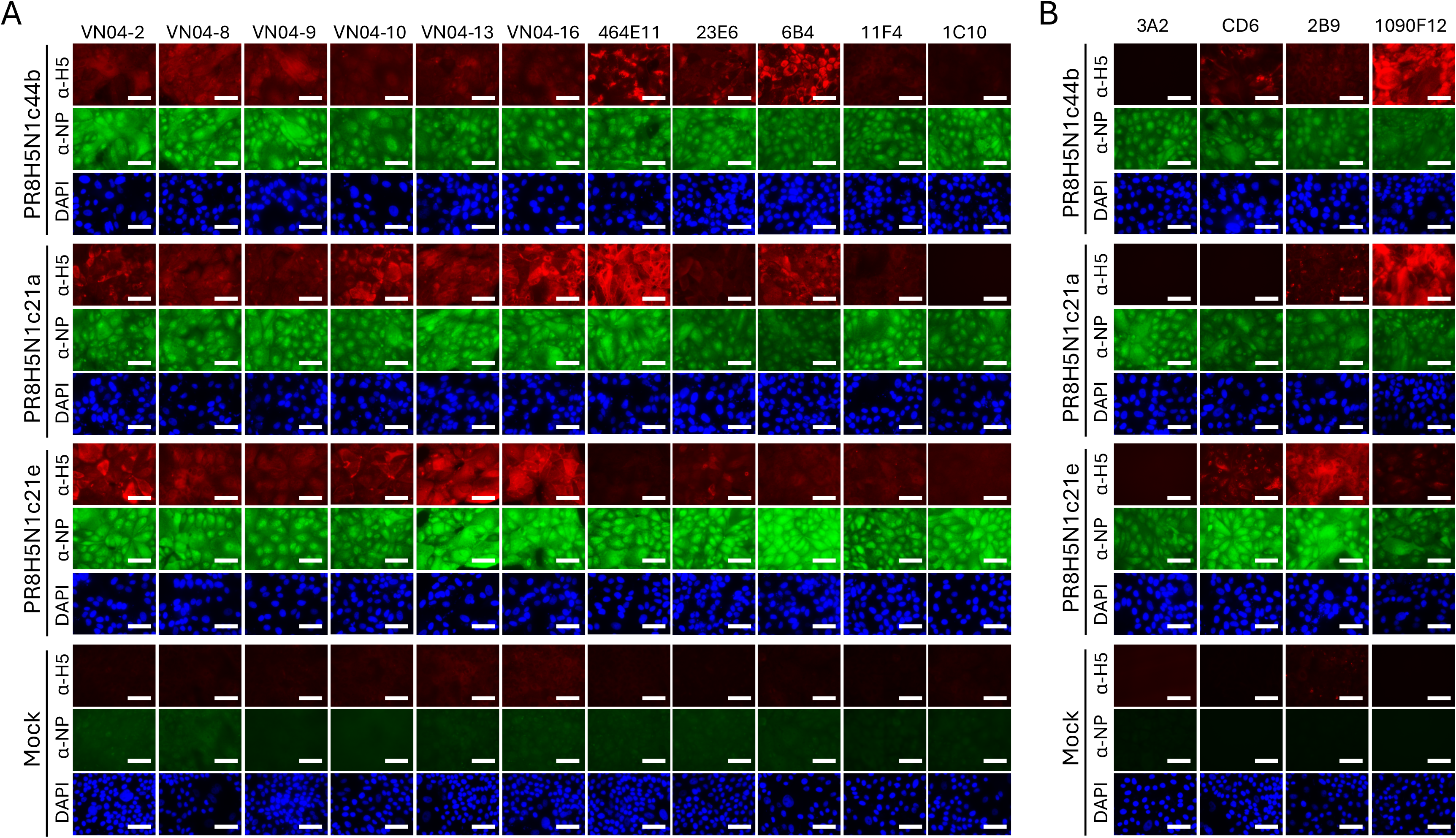
Cross-reactive recognition of anti-H5 and anti-N1 mAbs with the PR8-based viruses representing the three H5N1 clades by indirect immunofluorescence assay (IFA). (A) H5 mAbs: cross-reactivity of 11 anti-H5 mAbs with MDCK cells infected with PR8 viruses expressing the HA and NA glycoproteins of clade 2.3.4.4b (PR8H5N1c44b), clade 2.3.2.1a (PR8H5N1c21a), or clade 2.3.2.1e (PR8H5N1c21e) H5N1 viruses. (B) N1 mAbs: cross-reactivity of four anti-N1 mAbs with cells infected with the same PR8 viruses expressing the HA and NA glycoproteins of H5N1 viruses. Infected cells were co-immunostained with IAV rabbit polyclonal anti-NP Ab (green) to identify infected cells and the indicated anti-H5 (A) or anti-N1 (B) mAbs (red) to detect viral glycoprotein expression. Cell nuclei were counterstained with DAPI (blue). Mock-infected MDCK cells were used as controls for all the mAbs. Representative images are shown with scale bar of 50 μm.

**Figure 2.**
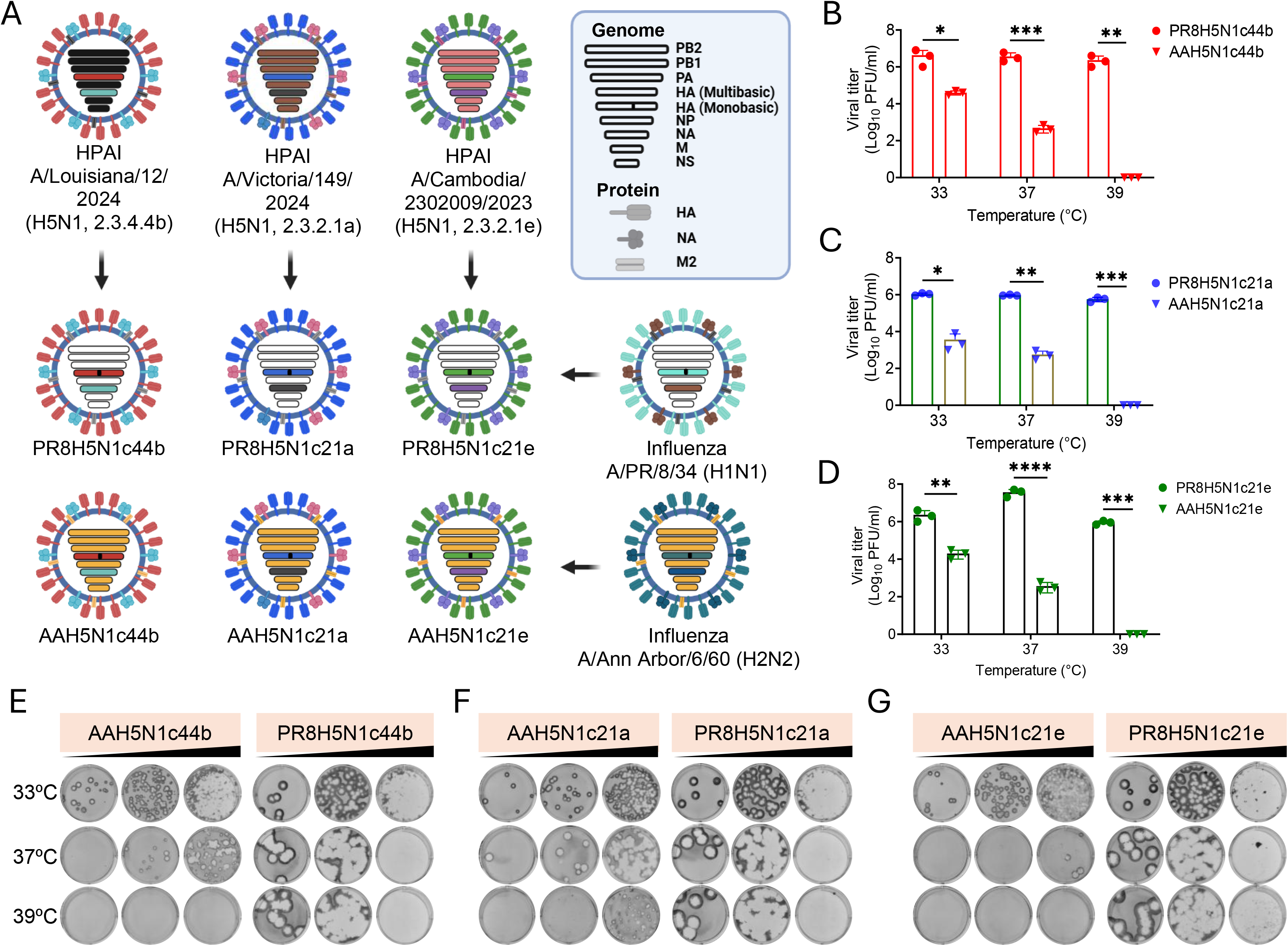
Generation and characterization of the AAH5N1c44b, AAH5N1c21a, and AAH5N1c21e viruses of the TriAAH5N1 LAIV. (A) Illustration of PR8– and AA-based H5N1 viruses. (B-D) Viral replication: Virus replication at 72 h post-infection (hpi) at different temperatures (33°C, 37°C, and 39°C) for PR8H5N1c44b and AAH5N1c44b (**B**), PR8H5N1c21a and AAH5N1c21a (**C**), and PR8H5N1c21e and AAH5N1c21e (**D**). **(E-G) Plaque assay:** Plaque phenotype of AA-based (left) and PR8-based (right) viruses at 72 hpi at the indicated temperatures (33°C, 37°C, and 39°C). Statistical analyses were performed using GraphPad Prism (v10.0, GraphPad Software). Comparisons involving multiple groups were analyzed using two-way analysis of variance (ANOVA) followed by Šídák’s multiple comparisons test. P values < 0.05 were considered statistically significant. Statistical significance is indicated as follows: ns, not significant; P < 0.05 (*); P < 0.01 (**); P < 0.001 (***); and P < 0.0001 (****).

To further define the antigenic relationships among these H5 clades, ferret antisera raised against candidate vaccine viruses (CVVs) A/chicken/Ghana/AVL-763_21VIR7050-39/2021 (IDCDC-RG80A; clade 2.3.4.4b), A/Victoria/149/2024 (NIID-003; clade 2.3.2.1a), and A/Cambodia/2302009/2023 (IDCDC-RG75A; clade 2.3.2.1e) were obtained from the CDC and evaluated by microneutralization against the corresponding low-pathogenicity PR8-based H5N1 viruses (**Table 1**). As expected, each antiserum efficiently neutralized its homologous virus, with MN titers of 10,240; 1,280; and 5,120 against PR8H5N1c44b, PR8H5N1c21a, and PR8H5N1c21e, respectively (**Table 1**). However, the cross-neutralization differed substantially among the three antisera. Antisera raised against clade 2.3.4.4b and clade 2.3.2.1a CVVs (IDCDC-RG80A and NIID-003) showed limited heterologous neutralizing activity, with MN titers ranging from 20 to 80 against viruses from the other clades. In contrast, antiserum raised against the clade 2.3.2.1e CVV (IDCDC-RG75A) displayed broader heterologous neutralization, with MN titers of 160 against PR8H5N1c44b and 640 against the closely related PR8H5N1c21a, although these responses remained markedly lower than the homologous neutralization titer (MN = 5,120) (**Table 1**).In addition, IVIG preparations, which contain Abs primarily generated against circulating seasonal IAVs, exhibited high HI titers against H1N1 and H3N2 viruses but showed no detectable HI activity against PR8H5N1c44b, PR8H5N1c21a, and PR8H5N1c21e (<10) (**Table 2**). In contrast, MN assay revealed differential neutralizing activity against PR8H5N1c44b, PR8H5N1c21a, and PR8H5N1c21e, with MN titers of 80, 20, and <10, respectively. This observed neutralization capacity of IVIG is unlikely to be mediated by HA-specific Abs and is most plausibly mediated by NA-specific Abs, indicating antigenic differences among the NA proteins of the three H5N1 strains. Together, the differential binding profiles of the anti-H5 and anti-N1 mAbs and the asymmetric cross-neutralization observed with CVV ferret antisera and IVIG demonstrate that the three representative H5 clades are antigenically distinct while retaining partially conserved epitopes. These findings support the generation and evaluation of a multivalent vaccine strategy incorporating representative H5 and N1 antigens from clades 2.3.4.4b, 2.3.2.1a, and 2.3.2.1e to broaden immune recognition across currently circulating H5N1 viruses.

**Table 1.**
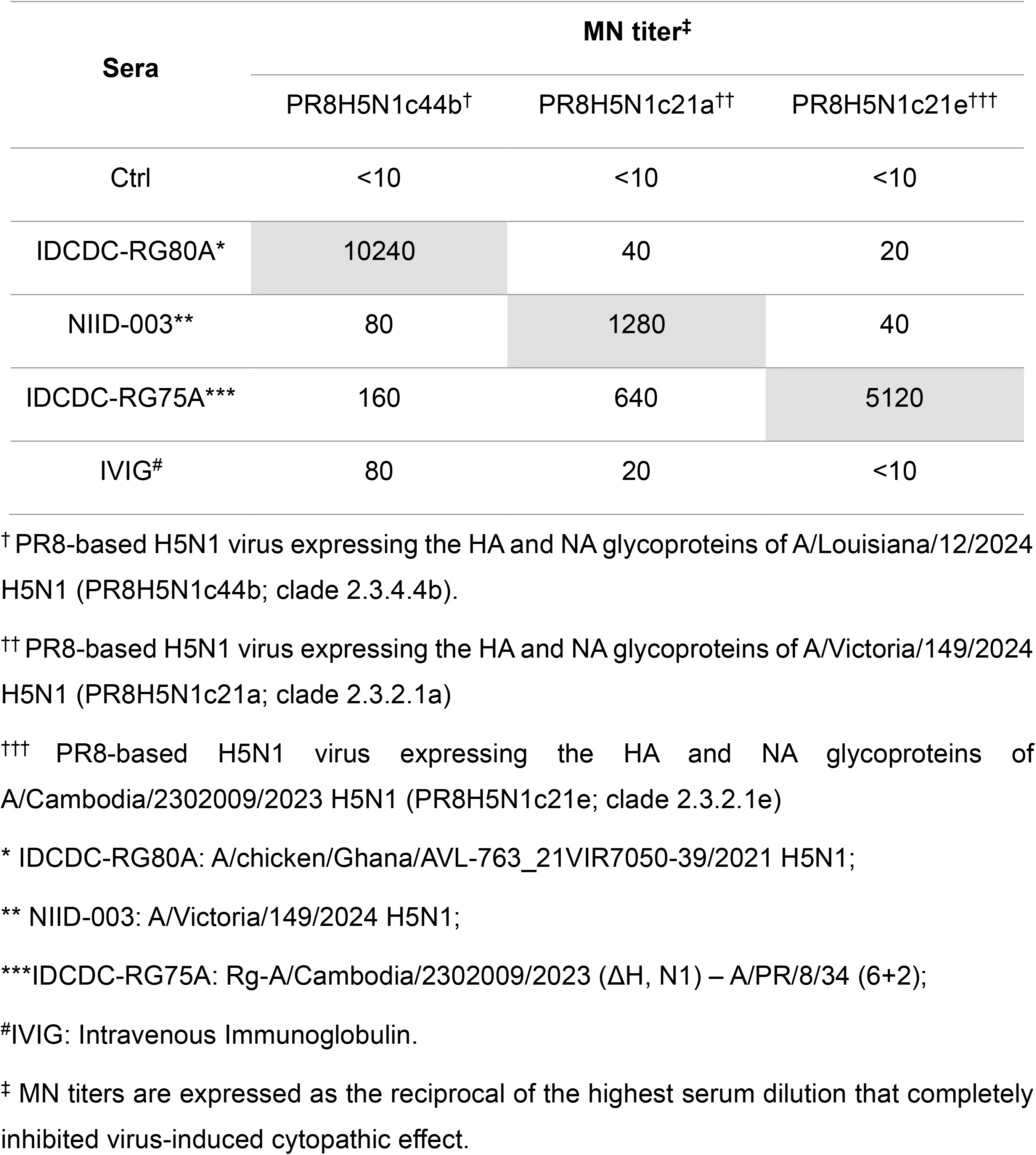
Cross-neutralization of representative of H5N1 clades virus by clade-specific ferret antisera and human intravenous immunoglobulin (IVIG).

**Table 2.**
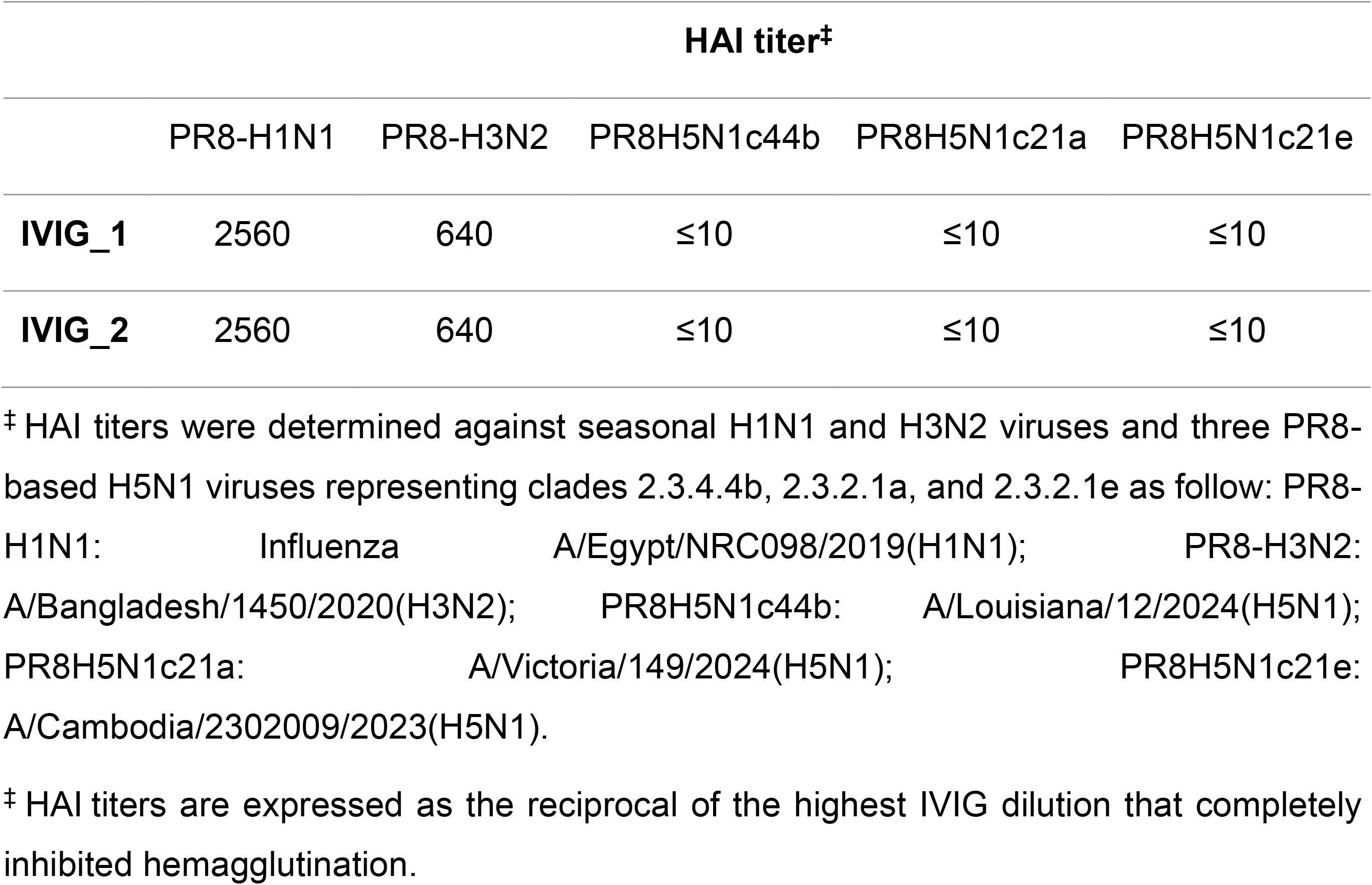
HAI activity titers of human IVIG against seasonal influenza viruses and divergent H5N1 clade viruses.

### Sequence analysis of HA and NA antigenic epitopes

To determine whether the observed antigenic differences were associated with variation in known H5 and N1 antigenic sites, the HA and NA sequences of the three representative H5N1 viruses were compared with the reference strain A/Vietnam/1203/2004 H5N1 (**Supplementary Tables S1; Supplementary Figs. S1 & S2**). All three viruses contained amino acid substitutions within established antigenic regions of the HA globular head, including residues located within or adjacent to the receptor-binding site (RBS), particularly the 130-, 150-, and 220-loop regions. Among the three viruses, clade 2.3.4.4b (A/Louisiana/12/2024) exhibited the highest number of substitutions within these antigenic regions, including Q115L, S123P, K140A, S155D, D183N, K189N, K218Q, and S223R. In comparison, clades 2.3.2.1a (A/Victoria/149/2024) and 2.3.2.1e (A/Cambodia/2302009/2023) retained more residues identical to the A/Vietnam/1203/2004 H5N1 HA, although each contained distinct clade-specific amino acid substitutions (**Supplementary Tables S1; Supplementary Figs. S1A and S1B**). Sequence analysis of N1 showed 81.7–89.8% amino acid identity with pH1N1 N1. Among the three H5N1 viruses, A/Louisiana/12/2024 (clade 2.3.4.4b) was the most closely related to pH1N1 (**Supplementary Fig. S2A**), retaining a full-length NA stalk, whereas A/Victoria/149/2024 and A/Cambodia/2302009/2023 contained a 20-amino acid stalk deletion. Unlike the HA, a few mutations in the NA antigenic epitopes have been observed (**Supplementary Tables S1 and Supplementary Figs. S2A-C**).

### Generation and *in vitro* characterization of the AA-based H5N1 multivalent LAIV candidate

Based on the antigenic and molecular characterization of the three representative H5N1 clades, we next generated, using reverse genetics, individual live-attenuated vaccine candidates using the influenza A/Ann Arbor/6/1960 master donor virus (MDV) backbone (**Fig. 2A**). To characterize their temperature-sensitive (*ts*) and cold-adapted (*ca*) phenotypes, the rescued vaccine viruses were compared with low-pathogenicity PR8-based H5N1 viruses at temperatures representing the human upper respiratory (33°C) and lower respiratory (37°C) track, and the febrile conditions associated with influenza infection (39°C). Plaque assays were developed in all cases at 72 hpi. As expected, the AA-based vaccine viruses replicated efficiently at 33°C but were markedly attenuated at 37°C and 39°C, demonstrating the characteristic *ts* and *ca* phenotype (**Figs. 2B-D**). Plaque morphology analysis further revealed significantly smaller plaques for the AA-based H5N1 viruses than for the PR8-based viruses, confirming their *ts* and *ca* phenotypes (**Figs. 2E-F**). In contrast, the PR8-based H5N1 viruses replicated efficiently and produced comparable plaque sizes at all tested temperatures (**Figs. 2B-2G**). These results demonstrate that the AA-based H5N1 viruses retain the characteristic *ts* phenotypes conferred by the AA MDV backbone, as evidenced by efficient replication at 33°C and limited or restricted replication with reduced plaque size at 37°C and 39°C.

### Safety and immunogenicity of the TriAAH5N1 LAIV in mice

Next, we evaluate the safety, immunogenicity, and protective efficacy of a trivalent AA-based H5N1 LAIV (TriAAH5N1) containing the three representative A/Louisiana/12/2024 (genotype D1.1, clade 2.3.4.4b; AAH5N1c44b), A/Victoria/149/2024 (clade 2.3.2.1a; AAH5N1c21a), and A/Cambodia/2302009/2023 (clade 2.3.2.1e; AAH5N1c21e) H5 and N1 proteins (**Fig. 3A**). These viruses were selected to represent genetically distinct circulating H5N1 clades, supporting the development of a broadly protective multiclade H5N1 vaccine candidate. Similarly, the three corresponding PR8-based H5N1 viruses were combined as a trivalent control (10^3^ PFU of each virus) and designated TriPR8H5N1. Untreated control (Ctrl) and mock-vaccinated (1XPBS; MVAC) mice were also included as control in these experiments (**Fig. 3B**). Body weight (**Fig. 3C**) and survival (**Fig. 3D**) in C57BL/6 mice were monitored daily for 14 days following intranasal (IN) inoculation, and mice reaching humane endpoints, defined as ≥25% body weight loss or severe clinical signs of infection, were humanely euthanized. Serum samples were collected weekly for four weeks (W1-W4) to assess vaccine-induced humoral immune responses. Remarkably, TriAAH5N1 LAIV was fully attenuated in C57BL/6 mice, with no detectable body weight loss or clinical signs of disease throughout the 14-day observation period, comparable to the Ctrl and MVAC groups (**Figs. 3C & 3D**). All TriAAH5N1-immunized mice survived until the end of the study, whereas infection with TriPR8H5N1 resulted in progressive body weight loss beginning at 2 days post-infection (DPI) and complete mortality between 6 and 7 DPI (**Figs. 3C & 3D**).

**Figure 3.**
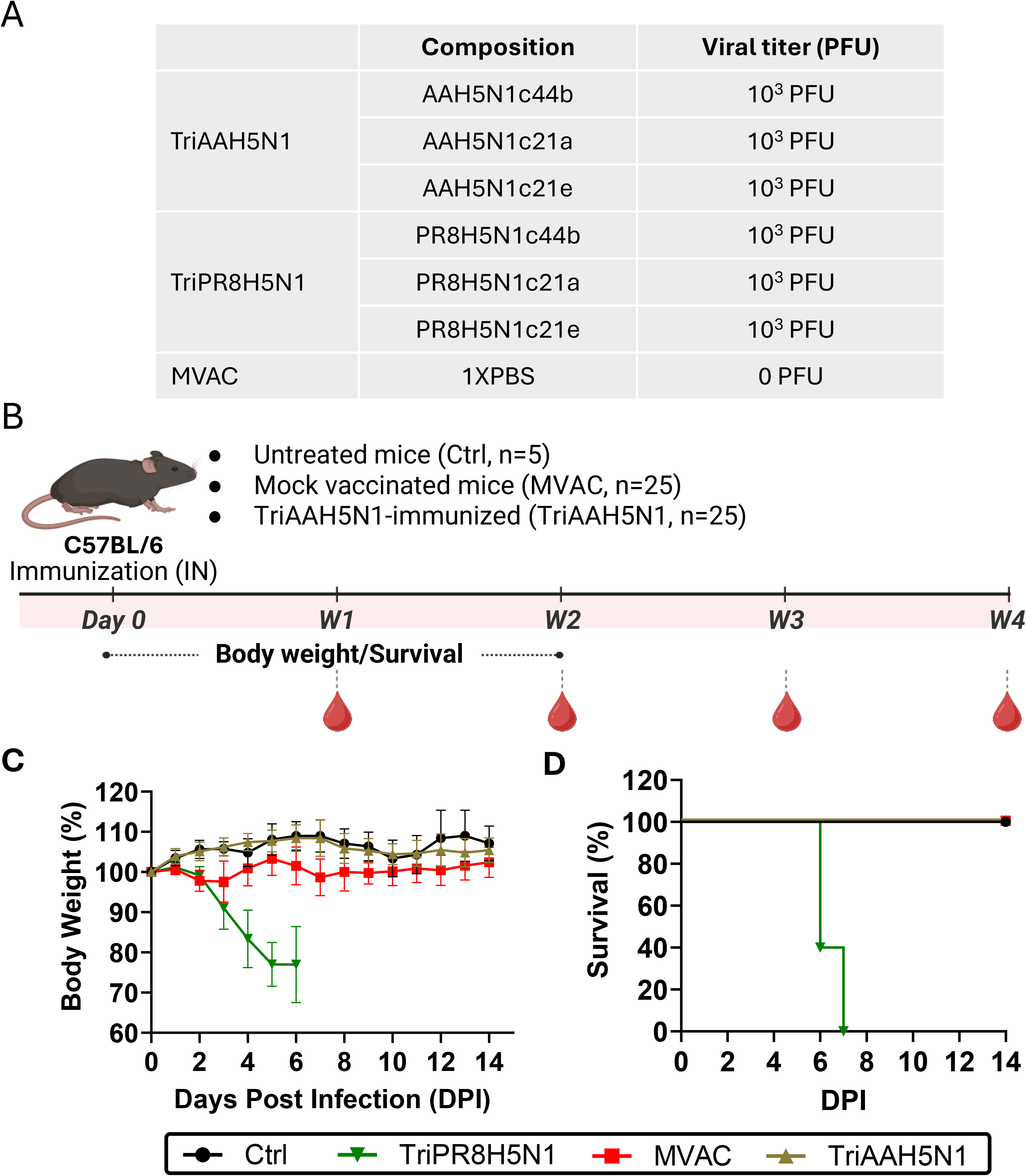
Safety of the TriAAH5N1 LAIV in C57BL/6 mice. (A) Viral composition: The composition of the TriAAH5N1 LAIV expressing the monobasic H5 HA and N1 NA glycoproteins from clade 2.3.4.4b (AAH5N1c44b), clade 2.3.2.1a (AAH5N1c21a), and clade 2.3.4.4e (AAH5N1c21e) H5N1 viruses. The three corresponding PR8-based H5N1 strains (PR8H5N1c44b, PR8H5N1c21a, and PR8H5N1c21e) were included as controls. MVAC: Mock-vaccinated control group. Ctrl: 1XPBS control group. **(B) Experimental design:** C57BL/6 mice were inoculated intranasally (IN) with the TriAAH5N1 LAIV (10^3^ PFU of each virus), TriPR8H5N1 (10^3^ PFU of each virus), or mock-vaccinated (1XPBS). Clinical signs of infection, changes in body weight, and mouse survival were monitored for 14 days post-infection (DPI). Sera samples were collected at weeks 1, 2, 3, and 4 after vaccination/infection. **(C) Body weight changes:** TriAAH5N1– and TriPR8H5N1-infected mice body weight was monitored and comparable to Ctrl and MVAC groups. **(D) Mortality:** Survival curves of TriAAH5N1– and TriPR8H5N1-infected and control C57BL/6 mice.

We next evaluated the immunogenicity and breadth of Ab cross-reactivity elicited by TriAAH5N1 LAIV against representative H5N1 viruses belonging to clades 2.3.2.1a, 2.3.2.1e, and the currently circulating clade 2.3.4.4b (D1.1 and B3.13 genotypes) (**Table 3**). Sera collected from TriAAH5N1-immunized mice exhibited robust neutralization activity against both clade 2.3.4.4b H5N1 viruses as early as one week after a single IN immunization, with neutralizing Ab titers ranging from 320 to ≥1,280. Importantly, these titers remained consistently high through four weeks post-immunization (W1-W4), indicating a sustained humoral immune response following a single vaccine dose. Similarly, sera from immunized mice demonstrated strong neutralizing activity against the clade 2.3.2.1e H5N1 virus, with neutralization titers of 640 at W1 that remained between 320 and 640 through W1-W4 post-immunization. Although the initial response against the antigenically distinct clade 2.3.2.1a H5N1 virus was lower (titer of 40 at W1), neutralizing titers increased progressively to 320 by W4 post-immunization, demonstrating continued maturation of the humoral immune response and increased Ab breadth over time. In contrast, sera collected from MVAC mice remained below the limit of detection (<10) against all H5N1 viruses tested throughout the study. Collectively, these findings demonstrate that a single IN immunization with the TriAAH5N1 LAIV is safe but able to induce rapid, sustained, and broadly cross-reactive neutralizing Ab responses against genetically identical and diverse H5N1 viruses, supporting its potential implementation as a multivalent pan-H5N1 LAIV.

**Table 3.**
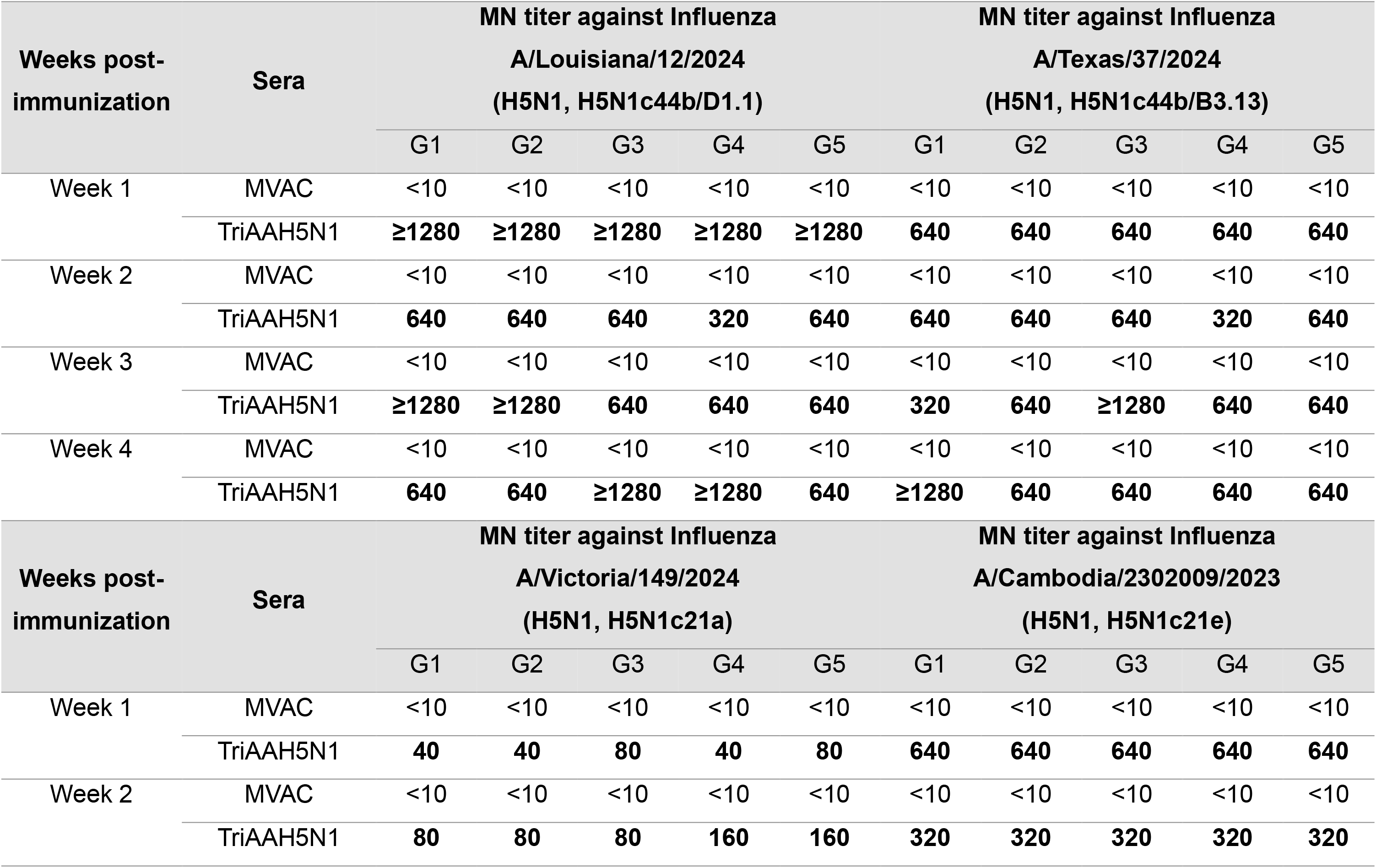
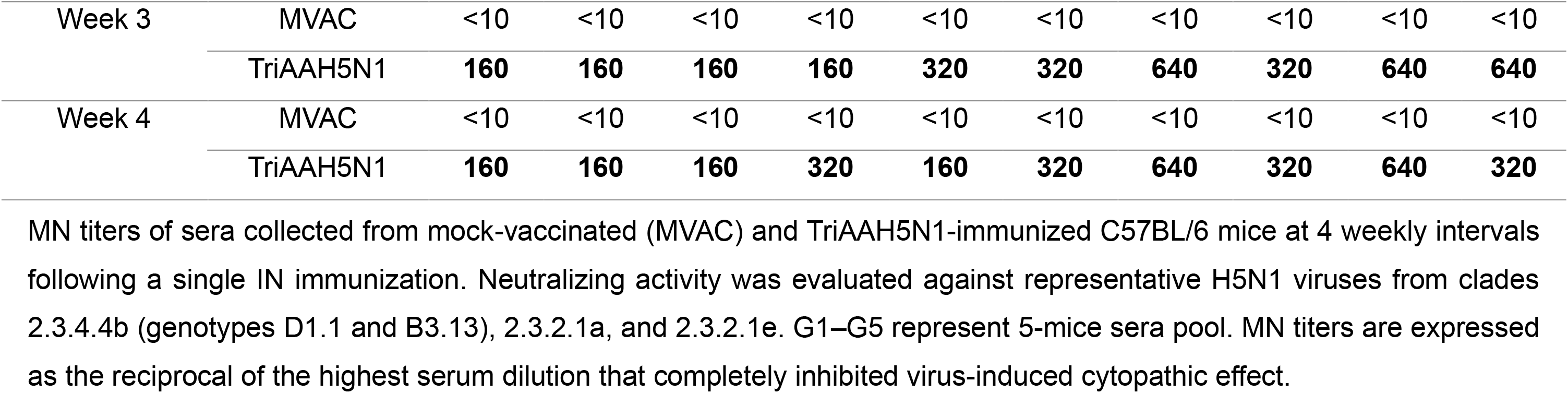
Longitudinal NAb responses induced by TriAAH5N1 IN vaccination against representative H5N1 viruses.

### Protective efficacy of the TriAAH5N1 LAIV against H5N1 viruses

To evaluate the protective efficacy of the TriAAH5N1 LAIV, C57BL/6 mice were IN immunized and subsequently challenged with 10^3^ PFU of H5N1 viruses expressing NanoLuc (Nluc), including influenza A/Texas/37/2024 H5N1 (HPhTX-Nluc), A/bovine/Texas/24-029328-02/2024 H5N1 (HPbTX-Nluc), A/Victoria/149/2024 H5N1 (HPhVIC-Nluc), and A/Cambodia/2302009/2023 H5N1 (HPhCAM-Nluc) (**Fig. 4A**). Control (Ctrl) and mock-vaccinated (MVAC) mice served as controls. Following challenge, clinical signs of infection, body weight, and survival were monitored daily for 14 days, and animals reaching the humane endpoint (≥25% body weight loss or severe clinical signs) were humanely euthanized. The TriAAH5N1 LAIV conferred complete protection against all challenge H5N1 viruses. Vaccinated mice maintained stable body weight throughout the observation period and exhibited 100% survival following challenge with each highly pathogenic H5N1 strain. In contrast, MVAC mice developed rapid weight loss and succumbed to infection between 5 and 7 DPI, with mortality occurring on day 5 following HPhTX-Nluc infection, day 6 following HPbTX-Nluc infection, and day 7 following HPhVIC-Nluc and HPhCAM-Nluc infections (**Fig. 4B**).

**Figure 4.**
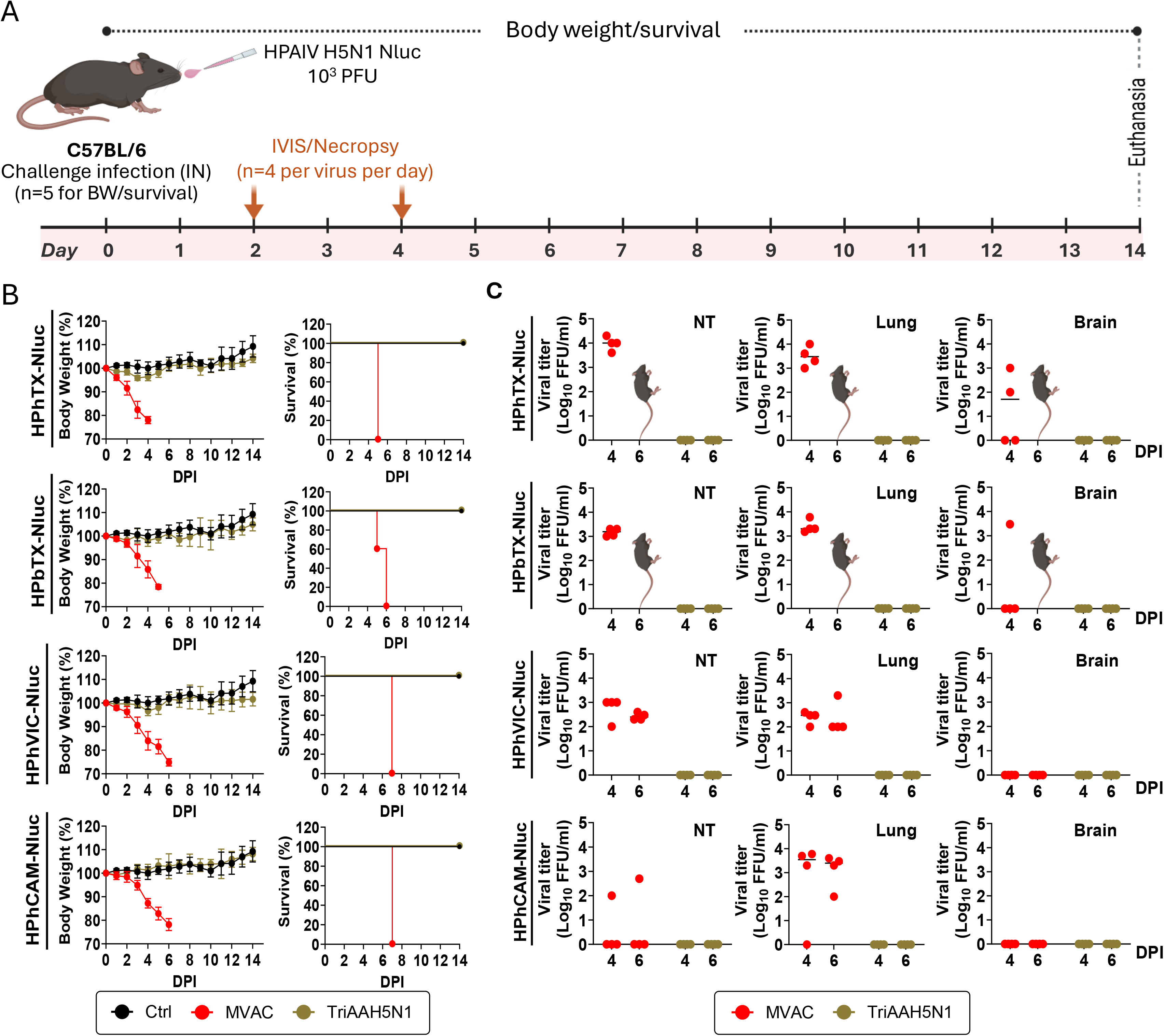
Protective efficacy of the TriAAH5N1 LAIV against challenge with divergent HPAI H5N1 clades. (A) Schematic representation of the study: C57BL/6 mice immunized IN with the TriAAH5N1 LAIV or mock-vaccinated (MVAC) were challenged with 10^3^ PFU of A/Texas/37/2024 (HPhTX-Nluc), A/bovine/Texas/24-029328-02/2024 (HPbTX-Nluc), A/Victoria/149/2024 (HPhVIC-Nluc), or A/Cambodia/2302009/2023 (HPhCAM-Nluc) expressing Nluc. Daily clinical observations, body weight changes, and survival were monitored for 14 days. At 4 and 6 DPI, IVIS and necropsy were conducted to evaluate Nluc expression and viral titers in different tissues. **(B) Body weight changes and survival rates following viral challenge:** Changes in body weight in C57BL/6 mice vaccinated with the TriAAH5N1 LAIV or mock-vaccinated (MVAC) after challenge with the indicated HPIV H5N1 viruses expressing Nluc. **(C) Viral loads:** Viral titers in nasal turbinate (NT), lungs, and brains at 4 and 6 DPI was evaluated by standard plaque assay in MDCK cells.

To assess viral replication, separate necropsy cohorts were euthanized on 4 and 6 DPI. Nasal turbinate (NT), lungs, and brains were collected for viral titration (**Fig. 4C**). High viral titers were detected in all examined tissues from MVAC mice, demonstrating extensive viral replication and systemic dissemination. In contrast, infectious challenge virus was undetectable in the NT, lungs, and brains of TriAAH5N1-immunized mice challenged with any of the four HPAI H5N1 viruses, indicating complete suppression of viral replication and efficient protection against viral challenge infection.

Protection was further validated by longitudinal *in vivo* bioluminescence imaging using Nluc reporter expression. Challenged mice were imaged using an *in vivo* imaging system (IVIS) prior to necropsy to monitor viral dissemination (**Fig. 5A**) followed by *ex vivo* imaging of the lungs (**Fig. 5B**). Strong Nluc signals were detected in all MVAC mice at both 4 and 6 DPI, confirming robust viral replication *in vivo*. *Ex vivo* imaging localized the bioluminescent Nluc signal predominantly to the lungs. In contrast, TriAAH5N1-immunized mice showed no detectable Nluc signal either *in vivo* or *ex vivo* following challenge with any of the recombinant Nluc-expressing H5N1 viruses, consistent with the complete absence of infectious virus detected by viral titration (**Fig. 4C**). Together, these findings demonstrate that the TriAAH5N1 LAIV provides complete protection against genetically diverse HPAI H5N1 viruses by preventing disease, eliminating viral replication in both the upper (NT) and lower (lungs) respiratory tracts, and blocking viral dissemination to peripheral tissues.

**Figure 5.**
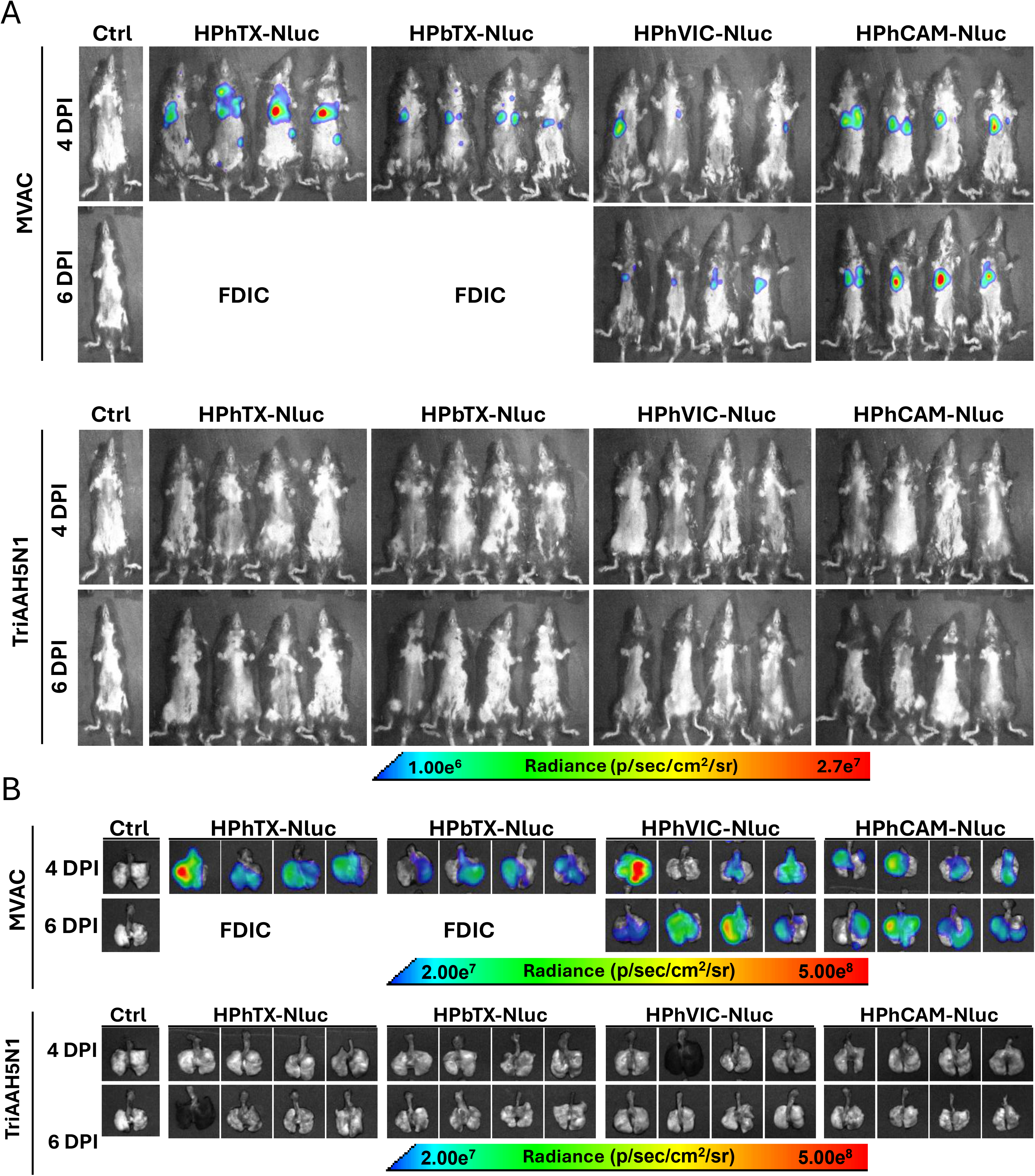
Protective efficacy of the TriAAH5N1 LAIV monitored by Nluc expression: Nluc expression was evaluated in TriAAH5N1-vaccinated and MVAC C57BL/6 mice at 4 and 6 DPI challenged with HPhTX-Nluc, HPbTX-Nluc, HPhVIC-Nluc and HPhCAM-Nluc (left to right) *in vivo* (**A**) and *ex vivo* (lungs) (**B**). FDIC: found dead in cage. Ctr: Control. Radiance, defined as the number of photons per s per square cm per steradian (ps^−1^ cm^−2^ sr^−1^), is shown on each of the indicated heatmaps.

### Impact of the TriAAH5N1 LAIV on virus-induced inflammatory innate immune responses

Severe AIV infection is characterized by an exaggerated innate immune response, often referred to as a cytokine storm, in which excessive production of pro-inflammatory cytokines and chemokines contributes substantially to lung pathology, acute respiratory distress, and mortality. Therefore, we evaluated the impact of TriAAH5N1 vaccination on mitigating excessive inflammatory cytokine and chemokine responses, including cytokine storm–associated mediators such as IFN-α, IFN-γ, IL-6, GM-CSF, CCL2, and CXCL10, following HPAI H5N1 infection (**Fig. 6**). Lung homogenates were collected from MVAC– and TriAAH5N1-vaccinated mice following challenge with HPAI H5N1 viruses representing clades 2.3.2.1a (HPhVIC), 2.3.2.1e (HPhCAM), and 2.3.4.4b (HPhTX). TriAAH5N1-vaccinated mice exhibited significantly lower levels of all measured inflammatory mediators than MVAC-vaccinated C57BL/6 mice (**Fig. 6**). Specifically, TriAAH5N1 vaccination significantly reduced the pulmonary production of IFN-α (**Fig. 6A**), IFN-γ (**Fig. 6B**), IL-6 (**Fig. 6C**), CCL2 (**Fig. 6D**), CXCL10 (**Fig. 6E**), and GM-CSF (**Fig. 6F**) following challenge with representative HPAI H5N1 strains compared to mock-vaccinate mice. These findings demonstrate that TriAAH5N1 vaccination not only limits viral disease but also markedly suppresses the excessive innate inflammatory response associated with HPAI H5N1 virus infections.

**Figure 6.**
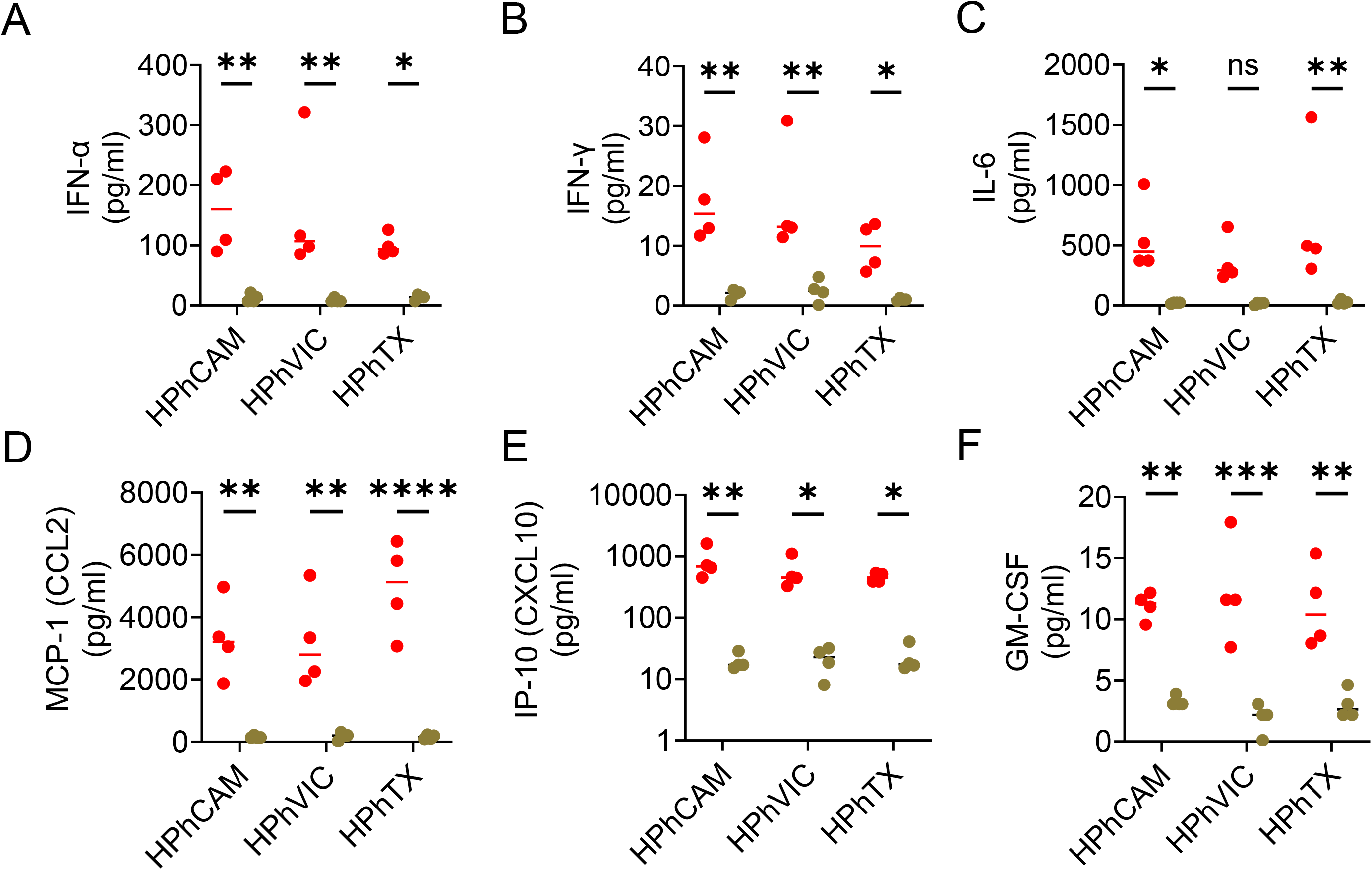
Vaccination with the TriAAH5N1 LAIV attenuates pro-inflammatory cytokine and chemokine responses following H5N1 viral challenge: Cytokine and chemokine levels were quantified using a multiplex Luminex assay in lung homogenates collected from control infected and vaccinated mice following challenge with clade 2.3.2.1a (HPhVIC), clade 2.3.2.1e (HPhCAM), and clade 2.3.4.4b (HPhTX) H5N1 viruses. The panel includes IFN-α (**A**), IFN-γ (**B**), IL-6 (**C**), CCL2 (**D**), CXCL10 (**E**), and GM-CSF (**F**). Data are presented as mean ± SEM. Statistical analyses were performed using GraphPad Prism (v10.5.0, GraphPad Software). Comparisons involving multiple groups were analyzed using two-way analysis of variance (ANOVA) followed by Tukey’s multiple comparisons test. P values < 0.05 were considered statistically significant. Statistical significance is indicated as follows: ns, not significant; P < 0.05 (*); P < 0.01 (**); P < 0.001 (***); and P < 0.0001 (****).

### Histopathological changes in lung tissues and detection of viral antigen by IHC

Histopathological changes and immunohistochemistry were also analyzed in lungs from mock-vaccinated and TriAAH5N1-vaccinated mice at 4 (**Figs. 7A-C**) and 6 (**Figs. 7D-F**) DPI with the H5N1 viruses. At 4 DPI (**Figs. 7A & 7B**), lungs from the mock control group show small BALT areas, absence of any infiltrative inflammatory cells, and absence of viral antigen staining (brown color) (**Fig. 7B**). Lungs from the mock-vaccinated and HPhTX challenged group showed an average 4 percent pathology, characterized by multifocal, moderate perivascular and peribronchiolar interstitial infiltration of lymphocytes, plasma cells and macrophages (**Fig. 7A**). The bronchial epithelium exhibited mild attenuation or sloughing with some karyorrhectic and cellular debris, infiltrating macrophages, lymphocytes, and plasma cells within the lumen. Immunohistochemistry for viral antigen showed diffuse, marked viral antigen staining (brown color) throughout the bronchial epithelium and various cells along and within the alveolar septa (staining average area 19.2% of the total lung surface) (**Figs. 7A & 7C**). In contrast, TriAAH5N1-vaccinated mice challenged with HPhTX only exhibited multifocal, mild BALT hyperplasia, with only scattered bronchial epithelial cells and few round cells within alveolar septa staining positive for influenza viral NP antigen (**Figs. 7B & 7C**). Mock-vaccinated mice challenged with HPhVIC showed an average 10.6 percent pathology characterized by multifocal areas of inflammation centered around or adjacent to bronchi and bronchioles. Affected bronchi and bronchioles had attenuated or sloughed epithelium with some karyorrhectic and cellular debris in the lumen, infiltrating macrophages, lymphocytes, and plasma cells. Alveolar septa surrounding the bronchioles were moderately thickened by infiltration of lymphocytes, plasma cells, macrophages. There was mild perivascular edema, and blood vessels were often lined by hypertrophied (reactive) endothelium. Immunohistochemistry for viral NP antigen showed diffuse, marked viral antigen staining throughout the bronchial epithelium and various cells along and within the alveolar septa (staining average 12.1% of the total lung surface). In contrast, 1 out of 4 TriAAH5N1-vaccinated mice challenged with HPhVIC, exhibited multifocal, mild BALT hyperplasia and 3 out of 4 mice showed multifocal, mild to moderate perivascular and peribronchiolar interstitial infiltration of lymphocytes, plasma cells and macrophages. Immunohistochemistry showed scattered bronchial epithelial cells and few round cells within alveolar septa staining positive for influenza NP viral antigen. One out of 4 mock-vaccinated mice challenged with HPhCAM showed mild inflammation (affecting 6% of the total lung surface) with alveolar septa surrounding the bronchioles thickened by infiltration of small numbers of lymphocytes, plasma cells, macrophages. Three out of 4 animals in the same group showed moderate inflammation, with lesions and diffuse viral antigen staining similar to the previous described MVAC and HPhVIC challenged group (group average lung pathology was 23% and viral NP staining average was 12.2% of the total lung surface). The TriAAH5N1-vaccinated mice challenged with HPhCAM only exhibited mild BALT hyperplasia multifocally and scattered bronchial epithelial cells and cells within alveolar septa stained positive for viral antigen.

**Figure 7.**
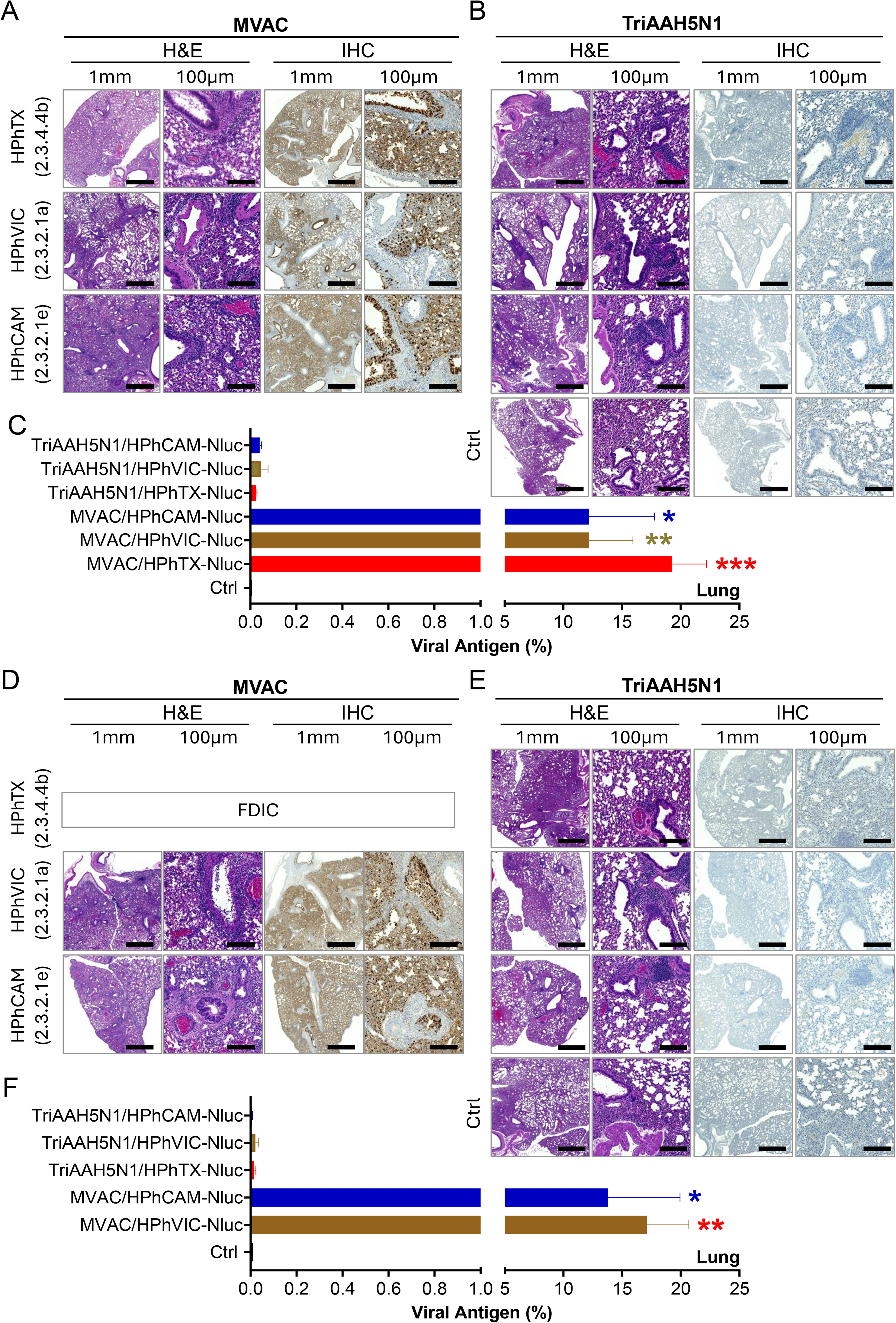
Histopathological and immunohistochemical analysis of lung tissues from MVAC– and TriAAH5N1-vaccinated mice following challenge with clades 2.3.4.4b, 2.3.2.1a, or 2.3.2.1e H5N1 viruses. (A-C) Lung tissues collected at 4 DPI: (**A**) Representative hematoxylin and eosin (H & E left) and immunohistochemistry (IHC; right) staining of lung sections from MVAC mice challenged with clades 2.3.4.4b, 2.3.2.1a, or 2.3.2.1e H5N1 viruses. (**B**) Representative H & E (left) and IHC (right) staining of lung sections from TriAAH5N1-vaccinated mice following challenge with the same H5N1 viruses. (**C**) Quantification of viral NP-positive areas in lung tissues from MVAC– and TriAAH5N1-vaccinated mice. **(D–F) Lung tissues collected at 6 DPI**: (**D**) Representative H&E (left) and IHC (right) staining of lung sections from MVAC mice following challenge with clades 2.3.4.4b, 2.3.2.1a, or 2.3.2.1e H5N1 viruses. (**E**) Representative H&E (left) and IHC (right) staining of lung sections from TriAAH5N1-vaccinated mice following challenge with the same H5N1 viruses. (**F**) Quantification of viral NP-positive areas in lung tissues from MVAC– and TriAAH5N1-vaccinated mice. Scale bars, 1 mm (low magnification) and 100 μm (high magnification). Statistical analyses were performed using GraphPad Prism (v10.5.0, GraphPad Software). Statistical analyses were done using an unpaired two-tailed Welch’s t-test. Statistical significance is indicated as follows: ns, not significant; P < 0.05 (*); P < 0.01 (**); P < 0.001 (***); and P < 0.0001 (****). Data are presented as mean ± SD. Differences in the percentage of NP-positive areas were analyzed using beta regression.

At 6 DPI (**Figs. 7D & 7E**), lungs from MVAC mice challenged with HPhTX show small BALT areas, absence of any infiltrative inflammatory cells, and absence of viral NP antigen staining (brown color). As described previously (survival curve), none of the mice mock vaccinated and challenge with HPhTX survived by 6 DPI. Both, mock-vaccinated mice challenged with HPhVIC and HPhCAM showed similar but increased severity in lung pathology compared to 4 DPI (the average percent lung pathology for HPhVIC challenged group was 30.6% and for HPhCAM group was 20.1%) (**Figs. 7D & 7F**). There was multifocal to coalescing large areas of inflammation centered around or adjacent to bronchi and bronchioles. Affected bronchi and bronchioles had attenuated or sloughed epithelium with karyorrhectic and cellular debris in the lumen, infiltrating macrophages, lymphocytes, and plasma cells. Alveolar septa surrounding the bronchioles are markedly thickened by infiltration of lymphocytes, plasma cells, macrophages. There was mild perivascular edema, and blood vessels were often lined by hypertrophied (reactive) endothelium. Immunohistochemistry for viral antigen showed marked, widespread viral antigen staining throughout the bronchial epithelium and cells within the alveolar septa that was greater than 4 DPI (the average percent lung viral antigen staining for HPhVIC challenged group was 17.1% and for HPhCAM challenged group was 13.8%) (**Figs. 7D & 7F**). The TriAAH5N1-vaccinated groups challenged with HPhTX and HPhCAM showed only multifocal, mild BALT hyperplasia and viral NP antigen IHC showed rare positive staining for viral antigen in scattered bronchial epithelial cells and few cells within alveolar septa. The TriAAH5N1-vaccinated group challenged with HPhVIC showed minimal inflammation with multifocal, moderate perivascular and peribronchiolar interstitial infiltration of lymphocytes, plasma cells and macrophages (average 2.3% lung pathology). IHC for viral NP antigen showed scattered positive staining in bronchial epithelial cells and few cells within alveolar septa (**Figs. 7E & 7F**). Altogether, these results demonstrate that TriAAH5N1 vaccination was associated with markedly reduced lung pathology and viral NP antigen detection following challenge with antigenically distinct HPAIV H5N1 strains at both 4 and 6 DPI.

### Protection efficacy of TriAAH5N1 LAIV against pH1N1

To assess the breadth of protection conferred by the TriAAH5N1 LAIV, vaccinated mice were challenged with the heterologous pH1N1 virus. This experiment was designed to determine whether immune responses elicited by the multivalent H5N1 LAIV extend beyond H5 viruses and provide cross-protection against antigenically distinct IAV sharing the N1 neuraminidase subtype present in currently circulating seasonal H1N1 IAV. To this point, MVAC– and TriAAH5N1-vaccinated mice were challenged with pH1N1-Nluc virus (10^3^ PFU/50µl/mice), and changes in body weight and survival were monitored for 14 days. MVAC mice began losing weight by 3 DPI (**Fig. 8A**) and all succumbed to infection between 7 and 8 DPI (**Fig. 8B**). In contrast, TriAAH5N1-vaccinated mice maintained their body weight throughout the 14 days observation period, exhibited no apparent clinical signs of disease, and all survived the challenge with pH1N1-Nluc (**Figs. 8A & 8B**). To evaluate viral dissemination, bioluminescence imaging (IVIS) was performed on a separate necropsy cohort at 6 DPI. *In vivo* imaging, followed by *ex vivo* imaging of the lungs, detected strong Nluc signal exclusively in mock-vaccinated animals, whereas no detectable Nluc signal was observed in TriAAH5N1-immunized mice (**Fig. 8C**). Viral titers were subsequently assessed in NT, lungs, and brain homogenates. High levels of pH1N1 replication were detected in MVAC mice, whereas TriAAH5N1-immunized animals showed significantly reduced viral titers in the NT and lungs, with virus levels at or below the limit of detection (LoD). As expected, no pH1N1 was detected in the brains of either group (**Fig. 8D**).

**Figure 8.**
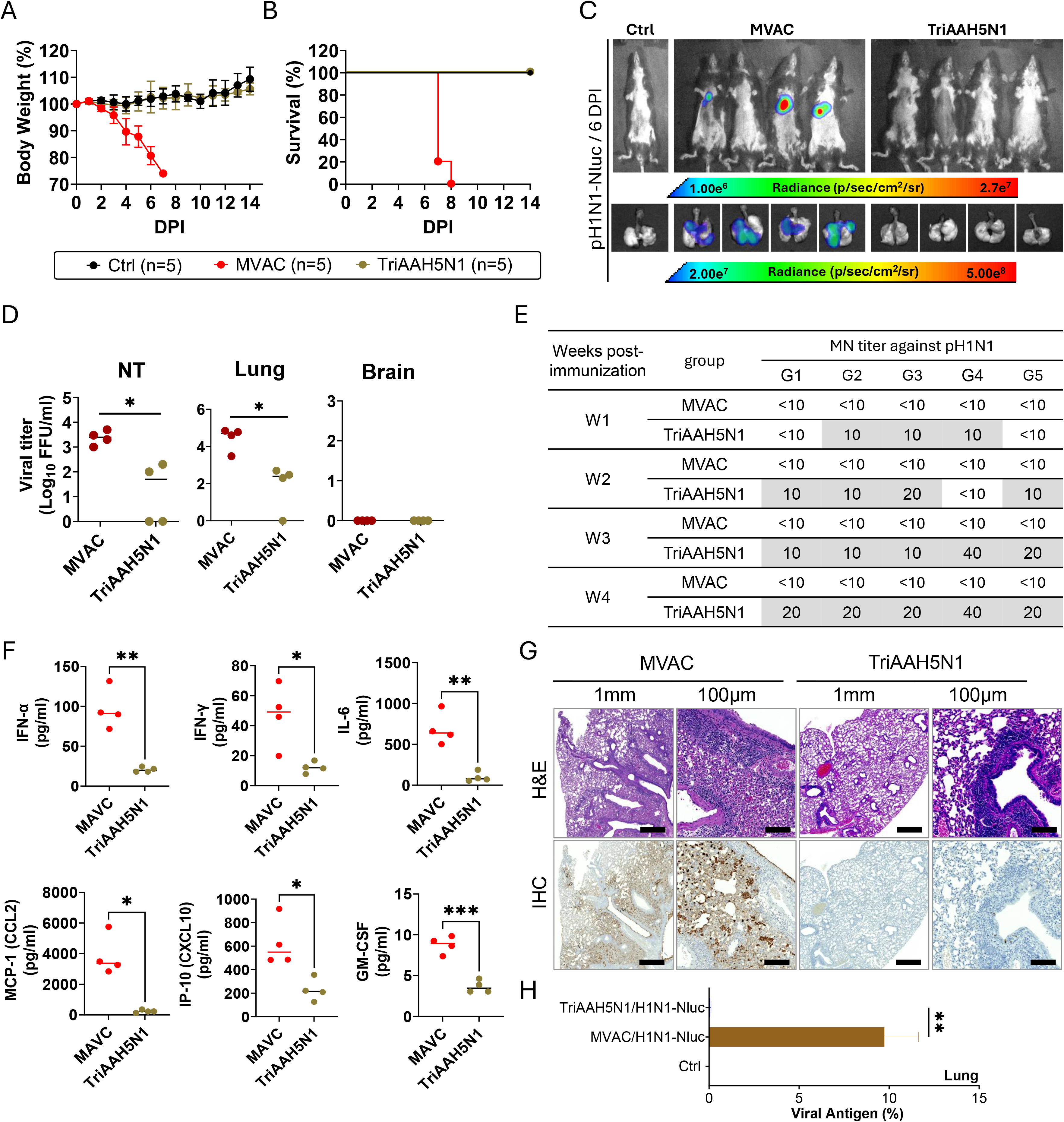
Cross-protective efficacy of the TriAAH5N1 LAIV against pandemic influenza A/California/04/2009 expressing Nluc (pH1N1-Nluc). (A-B) Clinical outcomes of TriAAH5N1-immunized and MVAC mice following challenge with pH1N1-Nluc. (A) Body weight changes following challenge. (B) Survival rates. (C) Representative *in vivo* bioluminescence images showing NanoLuc (Nluc) expression at the indicated DPI. Radiance (photons s⁻¹ cm⁻² sr⁻¹) is shown as a heat map. **(D) Viral titers of pH1N1-Nluc in NT, lungs, and brains (left to right) at 6 DPI determined by plaque assay on MDCK cells. (E) MN Ab titers against pH1N1 in sera collected at weeks 1–4 after immunization. (F) Lung cytokine and chemokine concentrations at 6 DPI determined by multiplex Luminex assay. (G) Representative hematoxylin and eosin (H & E left) and influenza NP immunohistochemistry (IHC; right) of lung sections collected at 6 DPI from MVAC– and TriAAH5N1-vaccinated mice following pH1N1-Nluc challenge.** Scale bars, 1 mm (low magnification) and 100 μm (high magnification). **(H) Quantification of viral NP antigen in the lungs of MVAC and TriAAH5N1-immunized mice.** Statistical analyses were performed using GraphPad Prism (v10.5.0, GraphPad Software). Statistical analyses were done using an unpaired two-tailed Welch’s t-test. Statistical significance is indicated as follows: ns, not significant; P < 0.05 (*); P < 0.01 (**); P < 0.001 (***); and P < 0.0001 (****). Data are presented as mean ± SEM. Statistical significance was determined using Mann-Whitney test, with p < 0.05 considered significant.

Because the TriAAH5N1 LAIV contains three H5 hemagglutinin antigens representing distinct clades together with three N1 neuraminidase antigens, we further evaluated the induction of cross-reactive Ab responses in TriAAH5N1-immunized mice sera against the heterologous pH1N1 virus. Microneutralization assays performed with sera collected before challenge demonstrated low but detectable neutralizing Ab titers (20–40) by weeks 3 and 4 post-vaccination, likely reflecting Abs directed against the conserved N1 NA shared between the TriAAH5N1 LAIV and the pH1N1 virus (**Fig. 8E**). The innate immune responses were also assessed in lung tissues collected after challenge. Interestingly, the lung homogenates of MVAC-mice exhibited markedly higher levels of pro-inflammatory cytokines and chemokines than TriAAH5N1-immunized animals, consistent with uncontrolled viral replication and increased pulmonary inflammation (**Fig. 8F**). Histopathological examination further supported these findings (**Figs. 8G & 8H**). At 6 DPI, MVAC mice challenged with pH1N1 showed moderate degree of inflammation (average percent pathology of 19.9%) characterized by multifocal areas of inflammation centered around or adjacent to bronchi and bronchioles. Affected bronchi and bronchioles had attenuated or sloughed epithelium with karyorrhectic and cellular debris in the lumen, infiltrating macrophages, lymphocytes, and plasma cells. Alveolar septa surrounding the bronchioles were thickened by infiltration of lymphocytes, plasma cells, macrophages. There was mild perivascular edema, and blood vessels are often lined by hypertrophied (reactive) endothelium (**Fig. 8G**). There was marked viral antigen staining throughout the bronchial epithelium and cells within the alveolar septa. In contrast, TriAAH5N1-vaccinated mice challenged with pH1N1 only showed multifocal, mild BALT hyperplasia and rare bronchial epithelial cells and few cells within alveolar septa stained positive for viral antigen (**Figs. 8G & 8H**). These results further demonstrate that the TriAAH5N1 LAIV provided effective control of viral replication following challenge with pH1N1.

### Safety profile of the TriAAH5N1 LAIV in immunocompromised C57BL/6 SCID mice

To investigate the safety and the role of adaptive immunity in vaccine-induced protection of the TriAAH5N1 LAIV against challenge with HPAI H5N1, we use SCID mice. SCID mice carry a spontaneous mutation in the *Prkdc* gene that results in the absence of functional T and B lymphocytes, making them an established model for evaluating vaccine safety and for determining whether protection depends on adaptive immune responses. To evaluate the safety of the TriAAH5N1 LAIV, SCID mice were IN inoculated with 10^3^ PFU of the TriAAH5N1 LAIV, while control (Ctrl) and MVAC were used as controls (**Fig. 9A**). Serum samples were collected weekly (W1-W4) to assess immunogenicity. The body weight and survival rates were monitored for 28 DPI. Mice inoculated with the TriAAH5N1 LAIV maintained their initial weight loss and lack clinical signs of disease, or mortality, demonstrating that the TriAAH5N1 LAIV was well tolerated in SCID mice lacking T and B cell responses (**Figs. 9B & 9C**). Consistent with the absence of functional B and T lymphocytes, TriAAH5N1-vaccinated SCID mice failed to generate detectable adaptive immune responses (**Supplementary Table S2**). Following the safety phase, mice were challenged IN with 10^3^ PFU of the homologous TriPR8H5N1 and clinical signs of infection, changes in body weight and survival were monitored for an additional 14 days (**Fig. 9D**). As expected, TriAAH5N1-vaccinated groups exhibited rapid weight loss (**Fig. 9E**) and 100% mortality (**Fig. 9F**), indistinguishable from MVAC SCID mice challenged with TriPR8H5N1 (**Figs. 9E & 9F**). These findings demonstrate that although the TriAAH5N1 LAIV is safe in severely immunocompromised mice, protection against lethal influenza challenge requires functional adaptive B and T cell immunity. Therefore, although safe, the efficacy of the TriAAH5N1 LAIV is expected to be limited in individuals with profound T– and B-cell deficiencies.

**Figure 9.**
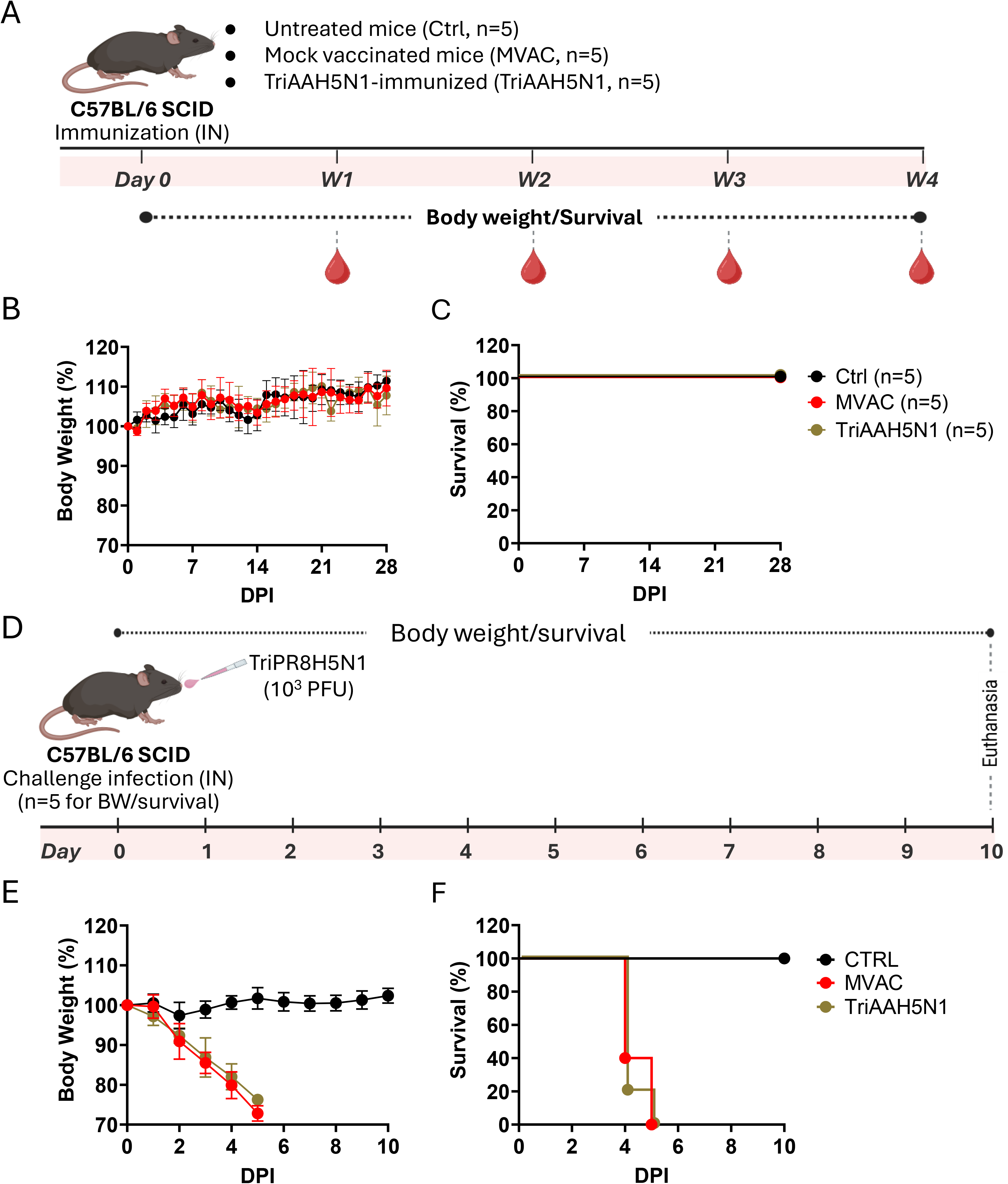
Safety and efficacy of the TriAAH5N1 LAIV in C57BL/6 SCID mice. (A-C) Vaccination with the TriAAH5N1 LAIV: C57BL/6 SCID mice, which lack functional T and B cells due to a spontaneous *Prkdc* mutation, were vaccinated IN with the TriAAH5N1 LAIV (10^3^ PFU of each clade) or mock-vaccinated (MVAC). Sera from vaccinated animals was collected at weeks 1, 2, 3, and 4 after vaccination (**A**) and body weight (**B**) and survival rates (**C**) were evaluated for 14 DPI. **(D–F) Challenge with TriPR8H5N1:** After week 4 post-vaccination with the TriAAH5N1 LAIV, SCID mice were challenge with TriPR8H5N1 (**D**) and body weight (**E**) and survival (**F**) were measured for 14 days (**D**).

## Discussion

Recent outbreaks of HPAI H5N1 clades 2.3.4.4b and 2.3.2.1 in mammals, including humans (30), have demonstrated the need for safe vaccines capable of inducing broad protection against multiple circulating variants rather than relying on strain-specific immunity. In this study, we developed a trivalent AA-based LAIV incorporating representative HA and NA antigens from three antigenically distinct HPAI H5N1 clades (TriAAH5N1). Our results demonstrate that this trivalent LAIV is highly attenuated, immunogenic, and capable of providing complete protection upon a single IN administration, against genetically diverse HPAI H5N1 viruses for it safe implementation as a pan-H5N1 vaccine.

The antigenic analyses confirmed that the selected H5N1 strains are distinct. Differential recognition by anti-H5 and anti-N1 mAbs, together with the asymmetric cross-neutralization observed with ferret antisera generated against CVVs demonstrated substantial antigenic divergence among clades 2.3.4.4b, 2.3.2.1a, and 2.3.2.1e H5N1 viruses despite the strong homologous neutralization. Sequence analysis provided a molecular explanation for these findings. The HA of A/Louisiana/12/2024 (clade 2.3.4.4b) contained the greatest number of substitutions within established antigenic regions, particularly the 140-loop and adjacent site B/Sa recognized by the VN04-series mAbs (31). Although these substitutions differ from the original escape mutations, they occur within the same Ab-binding regions and are consistent with the reduced recognition of the clade 2.3.4.4b virus by several A/Vietnam/1204/2004 H5N1 (VN04) mAbs. In contrast, the N1 proteins were considerably more conserved. Consistent with the sequence analysis, A/Louisiana/12/2024 H5N1, which retains a full-length NA stalk and is most closely related to pH1N1 N1, showed the highest IVIG-mediated neutralization, whereas the clade 2.3.2.1a and 2.3.2.1e H5N1 viruses exhibited reduced or absent neutralization. Together, these findings indicate that antigenic diversity exists in both HA and NA, although it is more pronounced in HA, supporting the inclusion of representative HA and NA antigens from multiple H5N1 clades in broadly protective vaccine formulations.

Interestingly, antisera against the clade 2.3.2.1e vaccine virus exhibited broader cross-neutralizing activity than antisera against the other two clades, suggesting that certain H5N1 viruses may preserve epitopes capable of inducing relatively broader immunity. This observation is consistent with a recent study showing that ferret antisera raised against clade 2.3.2.1e H5N1 cross-reacted with multiple contemporaneously circulating H5N1 clades, including clade 2.3.4.4b H5N1 viruses, as measured by HAI titers (32). In addition, an unexpected finding emerged from the analysis of IVIG. Although IVIG exhibited high HAI activity against seasonal H1N1 and H3N2 viruses, no detectable HAI activity was observed against any of the three HPAI H5N1 viruses. In contrast, low but measurable microneutralization activity was detected against PR8H5N1c44b and PR8H5N1c21a, but not against PR8H5N1c21e. Because HAI primarily measures Abs that bind and block the HA RBS, whereas virus neutralization can also be mediated by Abs targeting NA, these findings suggest that naturally acquired human immunity contains limited cross-reactive Abs against conserved N1 epitopes but little or no Ab reactivity to H5 HA (33, 34). The differences in neutralization among the three H5N1 viruses further indicate antigenic variation within the N1 despite their shared subtype designation. This interpretation is supported by a recent study showing that sera from individuals infected with pH1N1 exhibited little neuraminidase inhibition (NAI) activity against A/Bangladesh/Khulna/IEDCR-icddr,b-IC2/2025 clade 2.3.2.1a H5N1, whereas substantially higher NAI activity was detected against A/Texas/37/2024 clade 2.3.4.4b H5N1 (35). Together, these findings highlight the antigenic diversity of N1 proteins among H5N1 clades and support the inclusion of representative N1 antigens from multiple H5N1 clades in the design of broadly pan-H5N1 vaccines. Moreover, the antigenic analyses in this study provided a strong rationale for the design of the trivalent vaccine. Rather than selecting a single representative H5N1 strain, the TriAAH5N1 LAIV incorporates HA and NA antigens from three antigenically distinct H5N1 clades currently circulating in nature (13, 14, 16, 36). This allow broaden immune recognition and reduce the impact of antigenic mismatch caused by the rapid evolution of HPAI H5N1 viruses for the development and implementation of a pan-H5N1 vaccine.

The AA vaccine backbone retained the expected cold-adapted (*ca*), temperature-sensitive (*ts*), and attenuated (*att*) phenotypes after incorporation of the surface HA and NA glycoproteins of the three HPAI H5N1 viruses (37). Efficient replication at 33°C, together with marked attenuation at 37°C and 39°C of the three AA-based H5N1 viruses confirmed preservation of the established safety characteristics associated with licensed LAIV (37, 38). Consistent with these *in vitro* findings, the TriAAH5N1 LAIV was completely attenuated *in vivo* (37). Herein, mice vaccinated with the TriAAH5N1 LAIV exhibited no weight loss, no clinical signs of disease, and all survival infection, whereas mice infected with the PR8-based counterpart H5N1 viruses containing a monobasic cleavage site in their HA glycoproteins rapidly developed severe disease and succumbed to infection as previously described (39). These findings demonstrate that incorporation of multiple H5N1 antigens does not compromise the attenuation conferred by the AA backbone. Importantly, a single IN immunization with TriAAH5N1 LAIV generated rapid and sustained NAb responses against all three HPAI H5N1 clades. NAbs against clade 2.3.4.4b and 2.3.2.1e developed rapidly and remained high throughout the duration of the study (40, 41), while NAbs against the more antigenically distant H5N1 clade 2.3.2.1a increased progressively over time, suggesting continued affinity maturation and expansion of Ab breadth following vaccination. These data indicate that simultaneous presentation of multiple antigenically distinct HA molecules can induce broad humoral immunity after only one IN immunization. Importantly, the breadth of this immune response translated into complete protection against lethal challenge with four genetically distinct HPAI H5N1 viruses. Mice vaccinated with the TriAAH5N1 LAIV maintained body weight, all survived challenge with HPAI H5N1, and exhibited complete suppression of HPAI H5N1 replication in both the upper (NT) and lower (lung) respiratory tissues. The absence of detectable virus in the brain further demonstrates prevention of systemic viral dissemination, a hallmark of severe HPAI H5N1 infection. These findings were independently confirmed by *in vivo* and *ex vivo* bioluminescence imaging, which revealed complete absence of detectable viral replication in the TriAAH5N1-vaccinated animals.

Beyond preventing viral replication, TriAAH5N1 LAIV markedly reduced pulmonary inflammatory responses following challenge. Excessive production of cytokines and chemokines such as IL-6, IFN-α, IFN-γ, CXCL10, CCL2, and GM-CSF has been strongly associated with severe HPAI H5N1 disease and poor clinical outcomes. Mice vaccinated with the TriAAH5N1 LAIV exhibited significantly lower concentrations of each inflammatory mediator, consistent with rapid control of viral replication and prevention of a cytokine storm. Histopathological analysis further confirmed these observations, demonstrating minimal lung damage and little or no detectable viral NP antigen in TriAAH5N1-vaccinated animals compared with extensive pulmonary pathology in mock-vaccinated controls. A particularly interesting finding was the partial heterosubtypic protection against pH1N1. Although the TriAAH5N1 LAIV contains only H5 and N1 antigens, vaccinated mice were protected against a lethal challenge with pH1N1 and exhibited markedly reduced viral replication compared to mock-vaccinated mice. Because H5 and H1 HAs belong to different antigenic groups, this protection is unlikely to be mediated by HA-specific NAbs. Instead, the low but detectable MN titers observed against pH1N1 before challenge strongly suggest a contribution from cross-reactive N1 Abs, as both viruses shares the N1 subtype (22). In addition, LAIV are known to stimulate mucosal immunity, cellular immune responses, and trained innate immunity, all of which may contribute to broader heterologous protection (22). Consistent with our findings, a recent study reported that vaccination with a recombinant virus vaccine (rL H5) cross-protects against heterosubtypic H1N1 infection (42). In agreement with these results, several studies have demonstrated that pre-existing H1N1 immunity reduces disease severity caused by HPAI H5N1 infection (43, 44). Together, these findings suggest the potential for bidirectional cross-reactive Ab responses between seasonal and avian N1 NAs, whereby H5N1 vaccination or infection may induce Abs that recognize seasonal N1, and prior seasonal N1 immunity may similarly contribute to recognition of avian N1.

One important limitation of LAIV is that they are not recommended for immunocompromised individuals because of potential safety concerns (45, 46). The safety evaluation of the TriAAH5N1 LAIV in SCID mice further demonstrated that the vaccine itself remains attenuated even in immunocompromised hosts. TriAAH5N1-vaccinated SCID mice did not develop clinical disease, changes of body weight and all animals survive vaccination, indicating that the *att* phenotype of the TriAAH5N1 LAIV is stable in the absence of adaptive immunity. However, these animals failed to survive subsequent viral challenge with H5N1 because they were unable to generate antigen-specific immune responses. These findings confirm that TriAAH5N1 LAIV-mediated protection depends primarily on functional adaptive immunity rather than persistent viral attenuation alone.

Another limitation of LAIV is that they are only recommended to be used in individuals 2 through 49 years of age. The lower efficacy in older adults appears to be mediated by their limited replication due to excess levels of pre-existing immunity conferred by previous infections with seasonal IAVs. This limitation might be less relevant in the case of H5N1 vaccines due to the lower levels of pre-existing immunity in humans.

Clinical studies with monovalent H5N1 LAIV have been conducted in the past (47). Although in humans it appears that H5N1 LAIV are more attenuated and less replicative than seasonal LAIV, they were able to induce H5 specific Abs as measured by ELISA, although these Ab titers were not high enough to display HAI activity. This perhaps was due to the HA receptor specificity and/or the presence of N1 pre-existing immunity (48). Nevertheless, it was subsequently found that primed individuals with an H5 or H7 LAIV developed a rapid response to a booster with an IIV of the same subtype administered 3 months or 5 years later, characterized by the induction of high HAI titers (49, 50). Here we demonstrate that our generated TriAAH5N1 LAIV might expand even more protection against multiple antigenically diverse clades of H5N1 HPAIVs.

Altogether, our study demonstrates that a multiclade AA-based H5N1 LAIV can successfully overcome the antigenic diversity currently observed among circulating H5N1 viruses. By incorporating representative HA and NA antigens from three antigenically distinct H5N1 clades, the TriAAH5N1 LAIV induces broad NAb responses, completely protects against challenge with multiple HPAI H5N1 viruses, suppresses viral replication and inflammatory pathology, and provides cross-protection against pH1N1 in mice. These findings support further preclinical development of this multivalent IN H5N1 LAIV as a promising strategy to improve pandemic preparedness against the continuing evolution and risk of HPAI H5N1 viruses.

## Materials and Methods

### Biosafety

All experiments involving high pathogenicity and low pathogenicity avian influenza (HPAI and LPAI, respectively) H5N1 viruses were conducted in high containment biosafety level 3 (BSL3) and animal BSL3 (ABSL3) facilities at Texas Biomedical Research Institute (Texas Biomed). All procedures were conducted in accordance with protocols approved by the Texas Biomed Institutional Biosafety Committee (IBC #21-017) and Institutional Animal Care and Use Committee (IACUC # 1785 MU).

### Cells

Madin-Darby canine kidney (MDCK) and human embryonic kidney (HEK293T) cells were used to generate recombinant viruses. MDCK cells were used to propagate and generate stocks of recombinant viruses. Cells were cultured in Dulbecco’s modified Eagle medium (DMEM) (Invitrogen, USA) supplemented with 10% fetal bovine serum (FBS) and 1% PSG (penicillin, 100 U/mL; streptomycin, 100 μg/mL; L-glutamine, 2 mM) at 37°C with 5% CO_2_.

### Sera and monoclonal antibodies (mAbs)

The ferret antisera produced against 2.3.4.4b candidate vaccine viruses (CVV) influenza A/Astrakhan/3212/2020-like (H5N8, clade 2.3.4.4b; IDCDC-RG71A), A/American Wigeon/South Carolina/22-000345-001/2021 (H5N1, clade 2.3.4.4b; IDCDC-RG78A) and A/chicken/Ghana/AVL-763_21VIR7050-39/2021 (H5N1, clade 2.3.4.4b; IDCDC-RG80A), were kindly provided by Drs. Han Di and Bin Zhou at the Center for Disease Control and Prevention (CDC), Atlanta, Georgia, USA.

The anti-H5 mAbs NR-2728 (VN04-2), NR-2731 (VN04-8), NR-2734 (VN04-9), NR-2737 (VN04-10), NR-2740 (VN04-13), and NR-2743 (VN04-16) generated against influenza A/Vietnam/1204/2004 H5N1 were obtained from BEI Resources. The IAV nucleoprotein (NP)-specific mouse mAb HT103 and the anti-H5 mAbs 464E11, 23E6, 6B4, 11F4, and 1C10 were kindly provided by Dr. García-Sastre Laboratory at Icahn School of Medicine at Mount Sinai. MAbs 3A2 and CD6 (Abcam, USA), anti-NP polyclonal Ab PA5-32242 and anti-H5 mAb 2B9 (Thermo Fisher Scientific, USA) were obtained from commercial sources. The anti-N1 mAb 1090F12 was provided by Dr. Kobie Laboratory at University of Alabama in Birmingham.

### Generation of recombinant H5N1 viruses

To generate the PR8-based H5N1 viruses or the AA-based H5N1 LAIV viruses carrying the temperature-sensitive (*ts*), cold-adapted (*ca*), and attenuated (*att*) mutations, eight-plasmid reverse genetics systems were used. Briefly, a 1:1 co-culture of MDCK and HEK293T cells was co-transfected with 1 μg each of six plasmids encoding the PB2, PB1, PA, NP, M, and NS gene segments of PR8 or AA together with plasmids encoding a monobasic-cleavage-site-modified HA and the NA segments of A/Louisiana/12/2024 H5N1 (clade 2.3.4.4b), A/Victoria/149/2024 H5N1 (clade 2.3.2.1a), or A/Cambodia/2302009/2023 H5N1 (clade 2.3.2.1e) viruses, as previously described (51). Following virus rescue, recombinant viruses were propagated in MDCK cells using infection medium consisting of DMEM supplemented with 0.3% bovine serum albumin (BSA), 1% penicillin-streptomycin-glutamine (PSG), and 1 μg/mL TPCK-treated trypsin at 33°C. Successful virus rescue was confirmed by hemagglutination (HA) assay. Nanoluciferase (Nluc)-expressing influenza A/Texas/37/2024 H5N1 (HPhTX-Nluc), A/bovine/Texas/24-029328-02/2024 H5N1 (HPbTX-Nluc), A/Victoria/149/2024 H5N1 (HPhVIC-Nluc), A/Cambodia/2302009/2023 H5N1 (HPhCAM-Nluc), and A/California/04/2009 pandemic H1N1 (pH1N1-Nluc) viruses were generated using eight plasmid reverse genetics systems and engineered to express Nluc from the NS gene segment, as previously described (52, 53).

### Plaque Assays and immunostaining

MDCK cells were seeded at a density of 1×10^6^ cells per well in 6-well plates and incubated overnight at 37°C in a humidified atmosphere containing 5% CO_2_. Confluent monolayers were infected with serial dilutions of virus for 1 h at 37 °C. Following viral adsorption, cells were overlaid with agar-containing post-infection medium (1X DMEM, 0.3% BSA, 1% PSG, 1 µg/mL TPCK-treated trypsin, and 1% agar) and incubated until plaque development. At 24–72 h post-infection (hpi), depending on the virus, MDCK cell monolayers were fixed overnight with 10% neutral-buffered formalin. For plaque visualization, monolayers were stained with 0.1% crystal violet solution for 5 min at room temperature (RT) and rinsed with water. For immunostaining, cells were permeabilized with 0.5% Triton X-100 in 1X Phosphate-Buffered Saline (1XPBS) solution for 15 min at RT, blocked with 2.5% BSA and then incubated with the NP-specific mouse mAb HT103 (1:100). Bound mAb was detected using the VECTASTAIN ABC kit (Vector Laboratories, USA) according to the manufacturer’s instructions. Plates were imaged using a ChemiDoc MP Imaging System (Bio-Rad, USA).

### Immunofluorescence assay (IFA)

MDCK cell monolayers were infected with recombinant PR8 viruses expressing the monobasic HA and NA glycoproteins of A/Louisiana/12/2024 H5N1 (PR8H5N1c44b), A/Victoria/149/2024 H5N1 (PR8H5N1c21a), or A/Cambodia/2302009/2023 H5N1 (PR8H5N1c21e) at a multiplicity of infection (MOI) of 3 for 1 h at 37 °C. Following viral adsorption, the viral inoculum was removed, infection medium (1X DMEM, 0.3% BSA, 1% PSG, 1 µg/mL TPCK-treated trypsin) was added, and cells were incubated for an additional 10 h at 37°C in a humidified incubator with 5% CO₂. After viral infection, cells were fixed with 10% neutral-buffered formalin for 1 h, permeabilized with 0.5% Triton X-100 in 1XPBS for 10 min, and blocked with 2.5% bovine serum albumin (BSA) for 30 min at RT. Cells were incubated with the indicated primary Abs (5 µg/ml of each anti-mouse H5 Ab and 1:1,000 dilution for the rabbit anti-NP polyclonal Ab PA5-32242 (Thermo Fisher Scientific, USA) for 1 h, followed by incubation with the appropriate fluorophore-conjugated secondary Abs (green-fluorescent Alexa Fluor® 488 goat anti-rabbit IgG Ab and red-fluorescent Alexa Fluor® 594 goat anti-mouse IgG Ab or Alexa Fluor® 594 goat anti-human IgG Ab) using the manufacturer’s recommended dilution. The cells were treated with DAPI (4’,6-diamidino-2-phenylindole) (Invitrogen, USA) to fluorescently label the cell nuclei. After extensive washing with 1XPBS, cells were maintained in 1XPBS until imaging using an EVOS M5000 Imaging System (Thermo Fisher Scientific, USA).

### Mouse experiments

Six-week-old female C57BL/6 wild-type (WT) and severe combined immunodeficiency disease (SCID) mice were obtained from The Jackson Laboratory and maintained under specific pathogen-free (SPF) conditions at the Texas Biomed animal facility. Mice were anesthetized by intraperitoneal (IP) administration of a ketamine/xylazine mixture and inoculated intranasally (IN) with 50 μL of virus inoculum diluted in 1XPBS to the indicated viral dose(s). For survival studies, body weight was recorded daily for 14 days, beginning on day of infection (day 0), and animals were monitored daily for clinical signs of disease, body weight changes, and survival. Mice reaching the predetermined humane endpoint, defined as ≥25% body weight loss or severe clinical signs, were humanely euthanized.

Survival was analyzed using Kaplan–Meier curves. For *in vivo* imaging, anesthetized mice received 100 μL of Nano-Glo luciferase substrate (Promega, USA) diluted 1:10 in 1XPBS by retro-orbital injection and were immediately imaged using a Spectrum *in vivo* imaging system (IVIS). Bioluminescence images were acquired and analyzed using Aura software (Spectral Instruments Imaging, USA). Following *in vivo* imaging, mice were humanly euthanized, and nasal turbinate (NT), lungs, and brains were collected. Lungs were subsequently imaged *ex vivo* using the Spectrum IVIS and were bisected. One half of the lung tissue was fixed in 10% neutral-buffered formalin for histopathological analysis and the remaining half homogenized in parallel with the NT and brain tissues using a Precellys tissue homogenizer. Tissue homogenate supernatants were used for viral titration by plaque assay/immunostaining, as well as for cytokine profiling using a Luminex assay, whereas formalin-fixed lung tissues were processed for histopathological and immunohistochemical analyses.

### Multiplex cytokine assays

Innate immune responses in lung tissues collected following viral challenge were evaluated by quantifying cytokine and chemokine levels. Lung tissue homogenate supernatants were diluted 1:2 in Universal Assay Buffer. Cytokine and chemokine concentrations, including IFN-α, IFN-γ, IL-6, TNF-α, MCP-1 (CCL2), CXCL2, IP-10 (CXCL10), and GM-CSF, were measured using a customized ProcartaPlex multiplex immunoassay (Thermo Fisher Scientific, USA) according to the manufacturer’s instructions. Following assay preparation, samples were decontaminated by overnight incubation in 1% paraformaldehyde (PFA) before removal from the ABSL3 facility. Plates were acquired on a Luminex 100/200 System (Thermo Fisher Scientific, USA) using xPONENT v4.3.309.1 with the following settings: gate 7,500–25,000, sample volume 50 μL, 50 beads per analyte, sample timeout of 60 s, and standard photomultiplier tube settings. Data were analyzed using xPONENT v4.3.309.1.

### Histopathology and immunohistochemistry (IHC)

The left lung lobes collected at necropsy were fixed in 10% neutral-buffered formalin, processed using a Tissue-Tek VIP tissue processor, dehydrated through graded ethanol, cleared in xylene, and embedded in paraffin using a ParaPro XLT embedding system as previously described (3). Paraffin blocks were sectioned at 4 μm using a Microm HM325 rotary microtome, mounted onto glass slides, and stained with H&E using a Varistain Gemini automated slide stainer according to standard protocols. Histopathological evaluation was performed by a board-certified veterinary pathologist. For IHC, 4 μm thick paraffin sections were mounted on positively charged slides and processed using a Discovery Ultra automated IHC/ISH staining platform (Roche, USA). Tissue sections were deparaffinized, subjected to antigen retrieval (cell conditioning), and endogenous peroxidase activity was blocked. IAV NP antigen was detected using the rabbit polyclonal Ab PA5-32242 (Thermo Fisher Scientific, USA, 1:1500 dilution) for 1 h at RT, followed by anti-rabbit HQ and anti-HQ HRP detection reagents. Immunoreactivity was visualized using ChromoMap DAB (Roche), and sections were counterstained with hematoxylin and bluing reagent. Whole-slide images were acquired using a Zeiss Axio Scan scanner and analyzed with HALO v4.0 software.

### Virus microneutralization (MN) assays

Neutralizing Ab responses were evaluated using a standard MN assay as previously described with minor modifications (54). Briefly, two-fold serial dilutions of mouse sera (heat inactivated for 30 min at 56°C) collected from vaccinated mice at weeks 1-4 post-vaccination were incubated with 100 plaque forming units (PFU) of the indicated wild-type (WT) viruses for 1 h at RT. The serum–virus mixtures were then added to confluent MDCK cell monolayers seeded in 96-well plates in quadruplicate. Following incubation at 37°C for 3-4 days, neutralizing activity was assessed by monitoring virus-induced cytopathic effect (CPE). Once CPE was observed in the virus-only control wells, cell monolayers were fixed with 4% PFA and stained with 0.1% crystal violet solution for 10 min. Neutralizing Ab titers were defined as the reciprocal of the highest serum dilution that completely prevented CPE.

### Hemagglutination inhibition (HAI) assays

To evaluate the cross-reactivity of HA-specific Abs present in two intravenous immunoglobulin (IVIG) preparations, HAI assays were performed as previously described (55). Briefly, the IVIG preparations were treated with receptor-destroying enzyme II (RDE-II; Denka Seiken, USA) for 20 h at 37°C, followed by heat inactivation at 56°C for 30 min. After treatment, 12.5 µL of each IVIG sample was added in triplicate to 96-well plates and mixed with an equal volume (12.5 µL) of diluted virus containing 4 hemagglutinating (HA) units. Following incubation for 30 min at RT, 25 µL of 1% turkey red blood cells (RBCs) were added to each well. After 45 min, the HAI titer was defined as the reciprocal of the highest IVIG dilution that completely inhibited hemagglutination.

### Sequence analysis and structural modeling

Sequences were assembled and aligned in Geneious Prime v2024.2 (MAFFT option) against the HA and NA reference sequences of influenza A/Vietnam/1204/2004 H5N1 and the pH1N1 NA sequence. Three-dimensional protein models were generated using SWISS-MODEL and AlphaFold3, validated by QMEAN and Ramachandran statistics, and visualized in UCSF ChimeraX v1.10.1. For SWISS-MODEL, templates were selected automatically, and the highest-ranking model for each sequence, based on GMQE and QMEAN scores, was retained. The experimentally determined structures of influenza A/Vietnam/1203/2004 H5N1 (HA, PDB: 6CFG; NA, PDB: 3CL2) served as reference structures for structural comparisons. Amino acid substitutions, relative to reference strains, were mapped onto the protein surfaces to identify antigenically and functionally relevant residues.

### Statistical analysis

Statistical analyses were performed using GraphPad Prism v10.5.0 (GraphPad Software, San Diego, CA, USA). Data are presented as mean ± standard deviation (SD) unless otherwise indicated. Comparisons between two groups were performed using an unpaired two-tailed Welch’s *t*-test. Comparisons involving multiple groups or time points were analyzed by two-way analysis of variance (ANOVA) followed by Šídák’s or Tukey’s multiple-comparison test, as appropriate. Histopathological lesion scores and viral antigen-positive areas were analyzed using beta regression, where appropriate. Differences were considered statistically significant at *P* < 0.05. Statistical significance is indicated as follows: ns, not significant; *P* < 0.05 (\**); P < 0.01 (**); P < 0.001 (\*\*\**); and *P* < 0.0001 (****).

## Declarations

### Authors Contributions

Conceptualization: A.M.E. and L.M-S.; Methodology: A.M.E., R.S.B., A.R., R.A.E., H.N., E.M.C., T.J., V.S., E.M.A. and A.N.; Data collection and interpretation: A.M.E, R.S.B., R.J.W., E.M.A., A.G-S and L.M-S.; Funding acquisition and resources: A.M.E., R.J.W., E.M.A., A.G-S., and L.M-S.; Writing—original draft preparation: A.M.E., V.S., and L.M-S.; Writing—review and editing: all authors have read and agreed to the published version of the manuscript.

## Acknowledgments

We thank Ariel Robles and Ashley Gay-Cobb at Texas Biomedical Research Institute for the technical assistance. We also thank the histology unit team staff, Dr. Renee Escalona and Mr. Colin Chuba, for their assistance in tissue staining/immunostaining experiments, and the Cell Biology Core Lab at Texas Biomed for assistance with the multiplex cytokine assay. We thank Drs. Han Di and Bin Zhou at the Center for Disease Control and Prevention (CDC) for providing the ferret antisera samples IDCDC-RG71A, IDCDC-RG78A, and IDCDC-RG80A. We also acknowledge BEI Resources for providing mAbs.

## Supplementary Figures

**Supplementary Figure S1.**
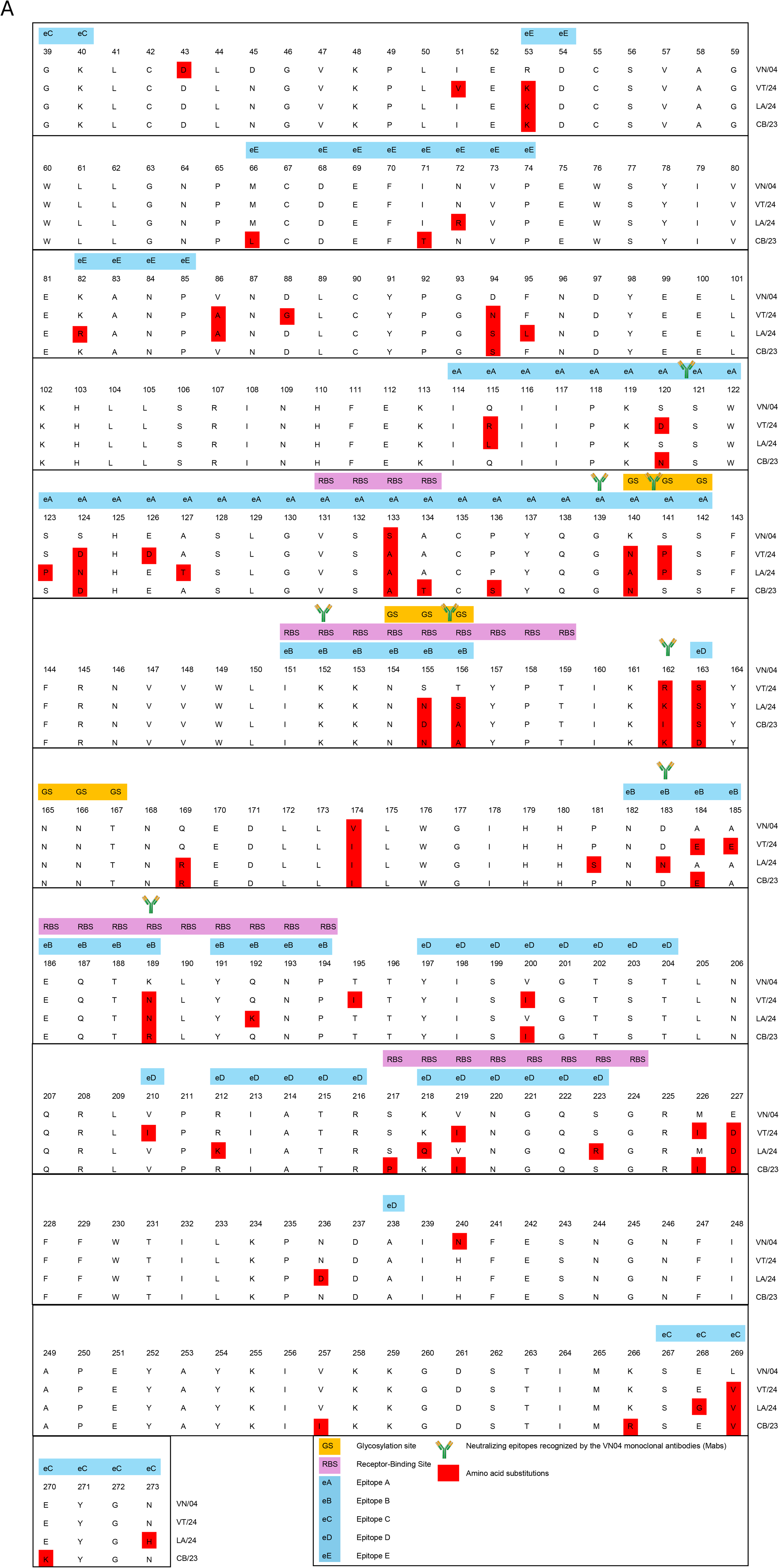
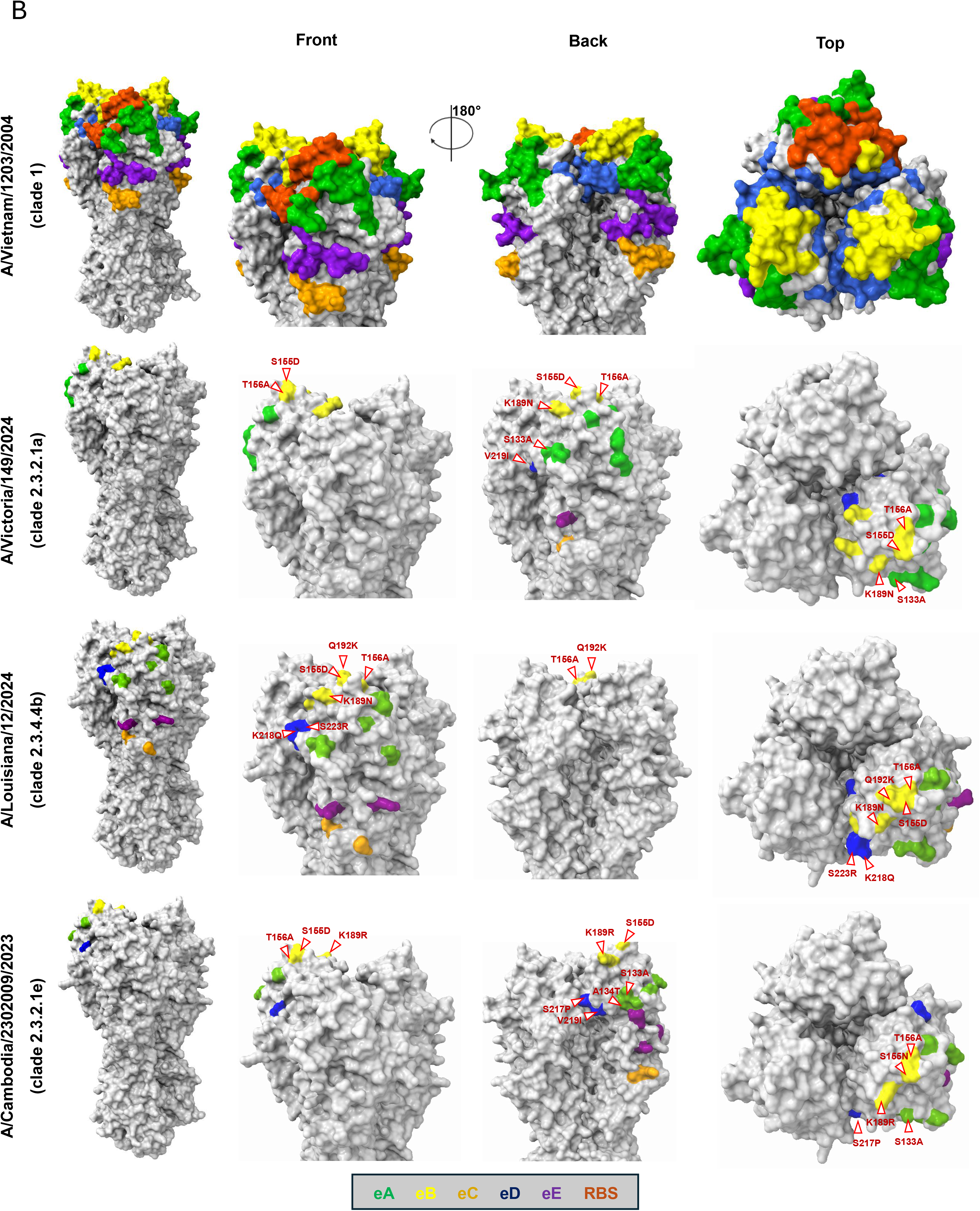
Comparative sequence and structural analysis of antigenic epitopes in H5 HA. (A) Sequence alignment of the five antigenic epitopes (eA-eE) of the H5 HA protein from the reference strain influenza A/Vietnam/1203/2004 (VN/04) H5N1 clade 1, and the three PR8-based H5N1 viruses VT/24 (A/Victoria/149/2024, clade 2.3.2.1a), LA/24 (A/Louisiana/12/2024, clade 2.3.4.4b), and CB/23 (A/Cambodia/2302009/2023, clade 2.3.2.1e). Only residues 39-273 are shown, encompassing all five antigenic epitopes (eA-eE). Residues recognized by VN/04 mAbs as reported by Kaverin et al (31), are indicated by an Ab icon. (**B) Front, side, and top views of the HA trimer showing amino acid substitutions relative to the reference strain A/Vietnam/1203/2004 (VN/04) H5N1 mapped onto the protein surface for each PR8-based H5N1 viruses:** Amino acid substitutions are shown in red and annotated according to H5 numbering. Antigenic epitopes are color-coded as follows: eA (residues 114-142, 131-134, green), eB (residues 151-156, 182-190, 192-195, yellow), eC (39-40, 267-273, orange), eD (163, 197-204, 210, 212-216, 218-223, 238, blue), and eE (53, 54, 66, 68-74, 82-85, purple). Residues 131-134 (130-loop), 151-159 (150-loop), 186-194 (190-helix), and 217-224 (220-loop), which comprise the RBS, are highlighted in maroon. Arrows indicate amino acid substitutions located within regions shared between antigenic epitopes and RBS. Three-dimensional models of the HA trimer were generated by SWISS Model using the amino acid sequence of A/Victoria/149/2024 H5N1, A/Louisiana/12/2024 H5N1, and A/Cambodia/2302009/2023 H5N1. The crystal structure of A/Vietnam/1203/2004 (VN/040 H5N1 (PDB: 6CFG) was used as the reference structure. Structural models were visualized and annotated using UCSF ChimeraX (version 1.10.1).

**Supplementary Figure S2.**
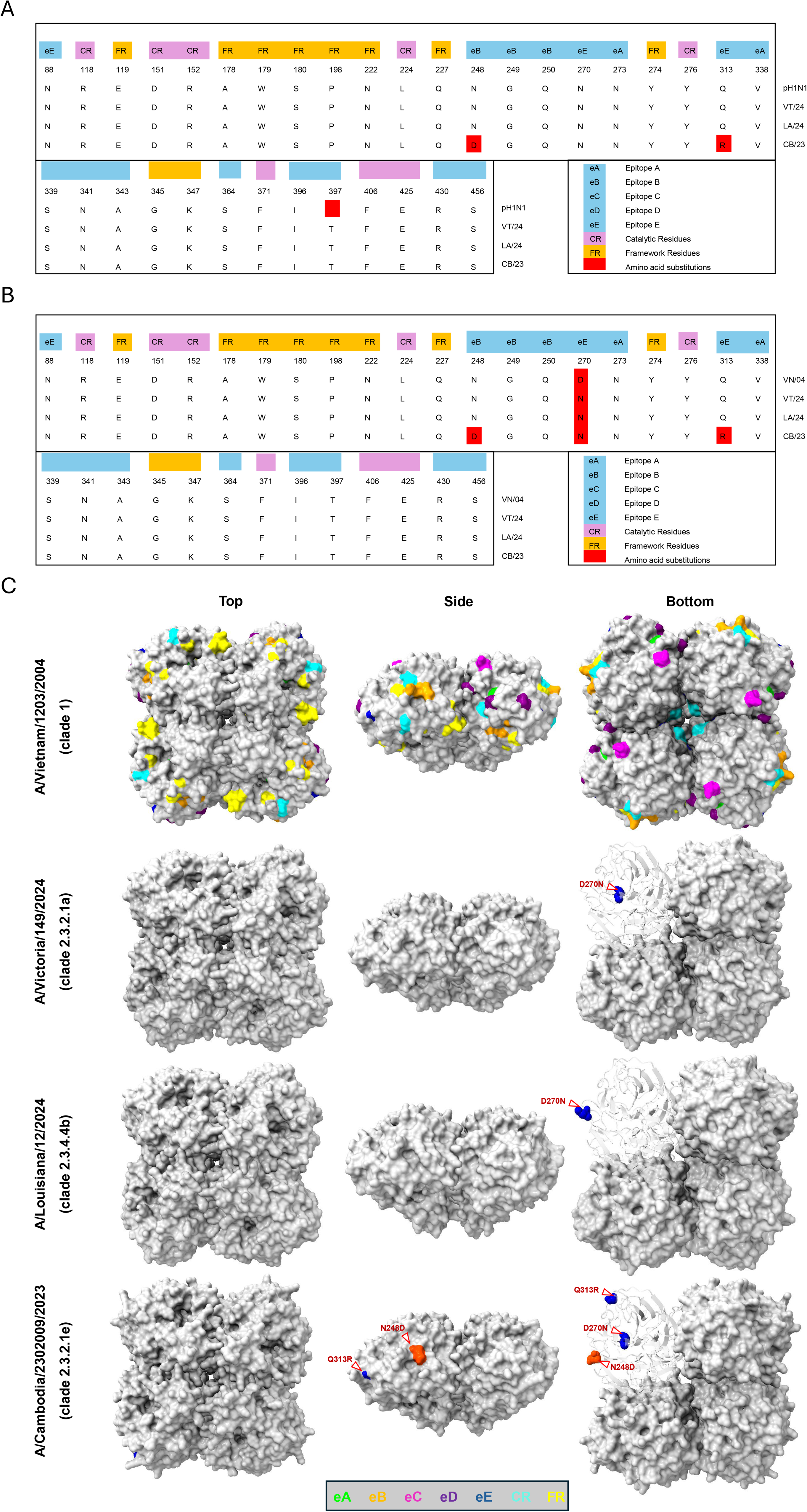
Sequence and structural comparison of the NA antigenic epitopes and catalytic residues. (A) Comparative sequence alignment of antigenic epitopes in the NA protein: Sequence alignment of the residues within the five antigenic epitopes (eA-eE), catalytic residues (CR; purple), and framework residues (FR; yellow) of the NA protein from reference strain A/California/04/2009 H1N1 (pH1N1) and the three PR8-based H5N1 viruses VT/24 (A/Victoria/149/2024 H5N1), LA/24 (A/Louisiana/12/2024 H5N1), and CB/23 (A/Cambodia/2302009/2023 H5N1). Amino acid substitutions relative to the pH1N1 reference are highlighted in red. **(B) Comparative sequence alignment of antigenic epitopes in the NA protein**: Sequence alignment of the residues within the five antigenic epitopes (eA-eE), catalytic residues (CR; purple), and framework residues (FR; yellow) of the NA protein from the reference H5N1 strain A/Vietnam/1203/2004 (VN04) and the three PR8-based H5N1 viruses VT/24 (A/Victoria/149/2024 H5N1), LA/24 (A/Louisiana/12/2024 H5N1), and CB/23 (A/Cambodia/2302009/2023 H5N1). Amino acid substitutions relative to the VN04 reference are highlighted in red. **(C) Top, side, and bottom views of the NA tetramer showing amino acid substitutions relative to the reference strain A/Vietnam/1203/2004 H5N1 (VN04) for the three PR8-based A/Victoria/149/2024 H5N1 (clade 2.3.2.1a), A/Louisiana/12/2024 H5N1 (clade 2.3.4.4b), and A/Cambodia/2302009/2023 H5N1 (clade 2.3.2.1e) viruses.** Substitutions are shown in red and numbered according to N1 numbering. Antigenic epitopes are color-coded: eA (lime; 273, 338, 339), eB (orange; 248, 249, 250, 341, 343), eC (magenta; 396, 397, 456), eD (purple; 364, 369, 430), and eE (blue; 88, 270, 313). Catalytic residues (CR) 118, 151,152, 224, 276, 292, 371, 406, 425 (N1 numbering) are shown in cyan and framework residues (FR) 119, 178, 179, 180, 198, 222, 227, 274, 294, 345, 347 in yellow. Arrows indicate substitutions located within or adjacent to antigenic epitopes or the catalytic/framework residues. Three-dimensional models of the NA tetramer were generated using SWISS Model using the amino acid sequence of A/Victoria/149/2024 H5N1 (VT/24, clade 2.3.2.1a), A/Louisiana/12/2024 H5N1 (LA/24, clade 2.3.4.4b), A/Cambodia/2302009/2023 H5N1 (CB/23, clade 2.3.2.1e) and using the amino acid sequence of influenza A/Vietnam/1203/2004 H5N1 (VN04) (PDB number 3CL2). Structural models were visualized and annotated using UCSF ChimeraX (version 1.10.1).

---

**Supplementary Table S1.**
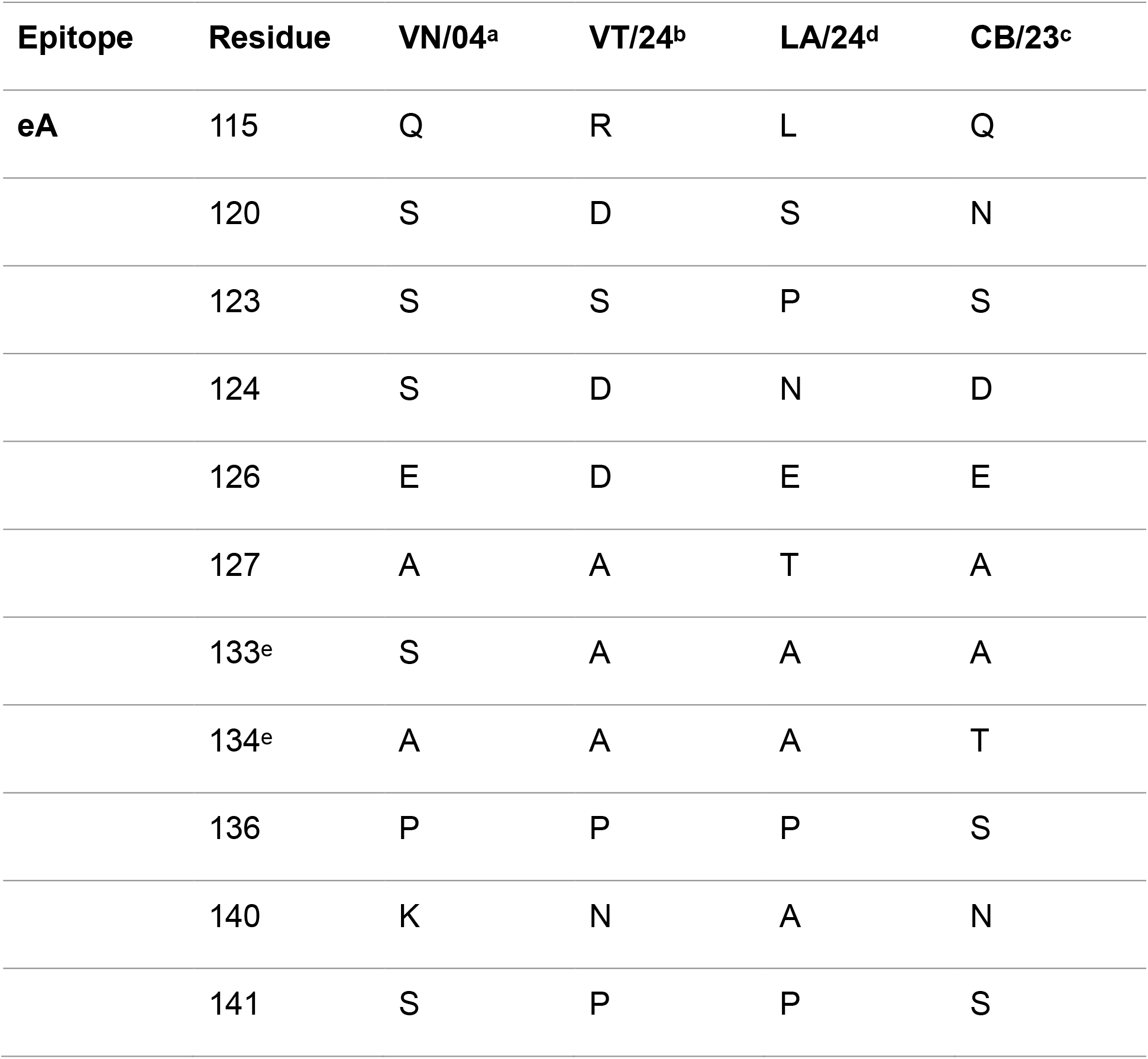
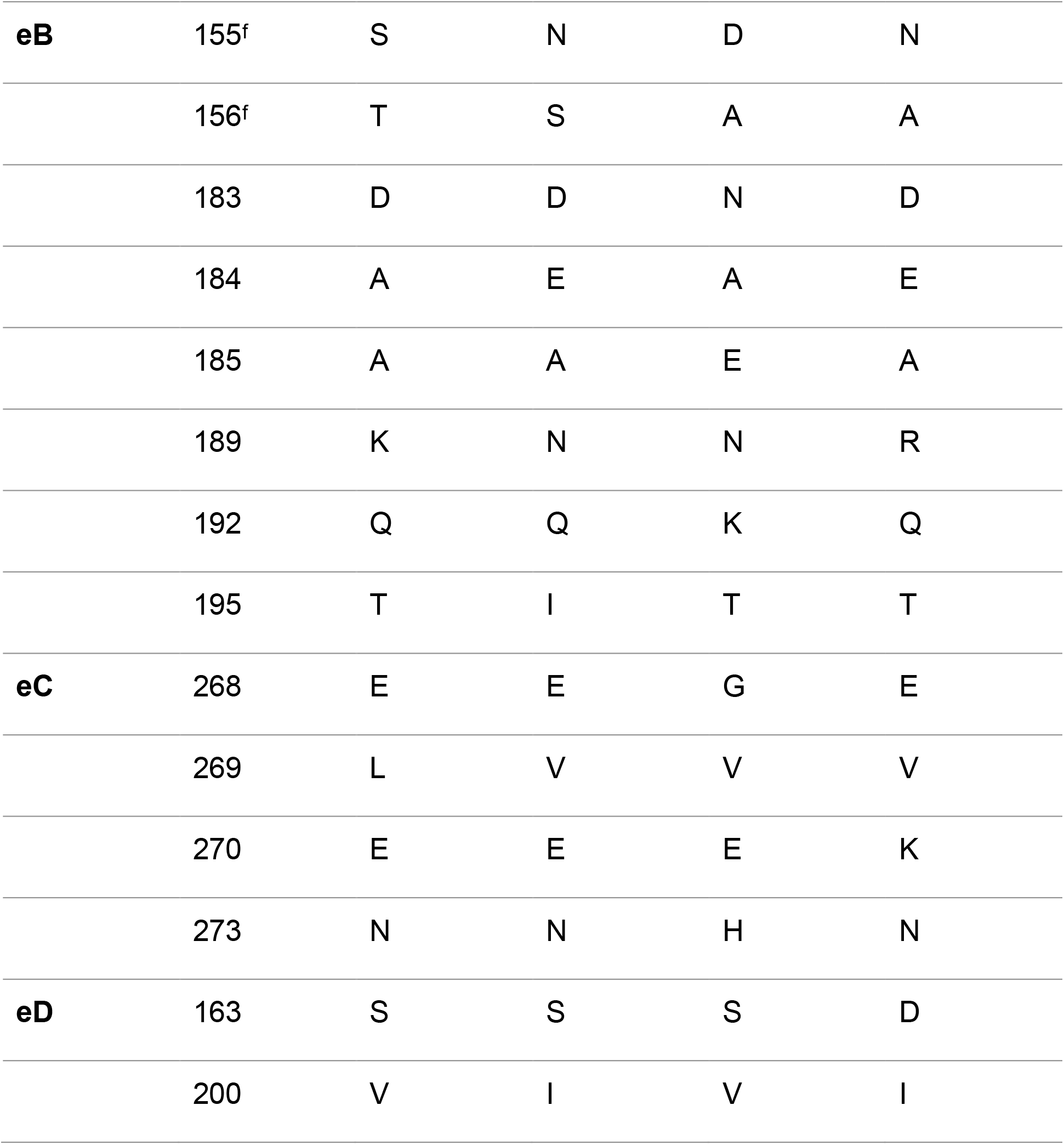
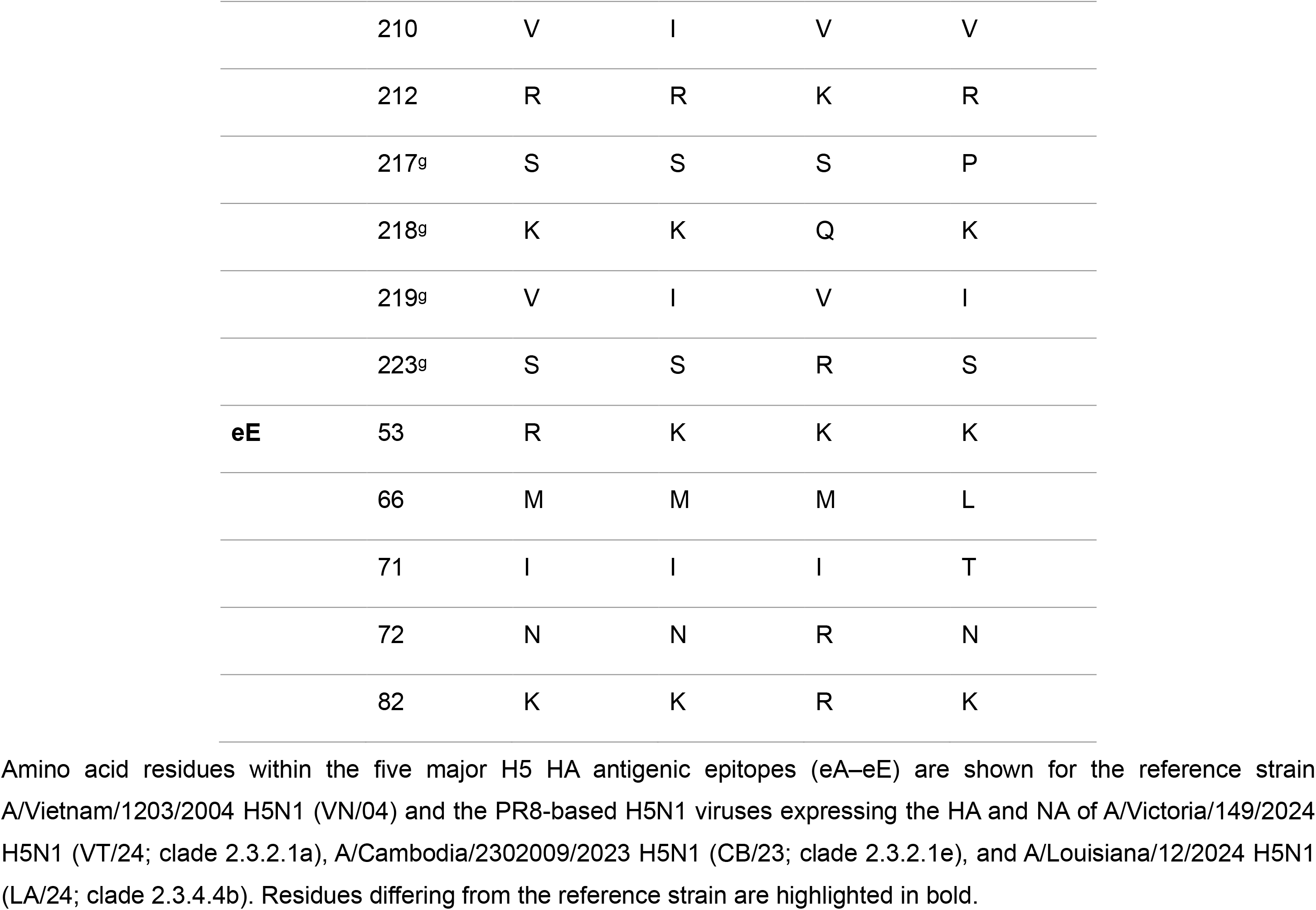
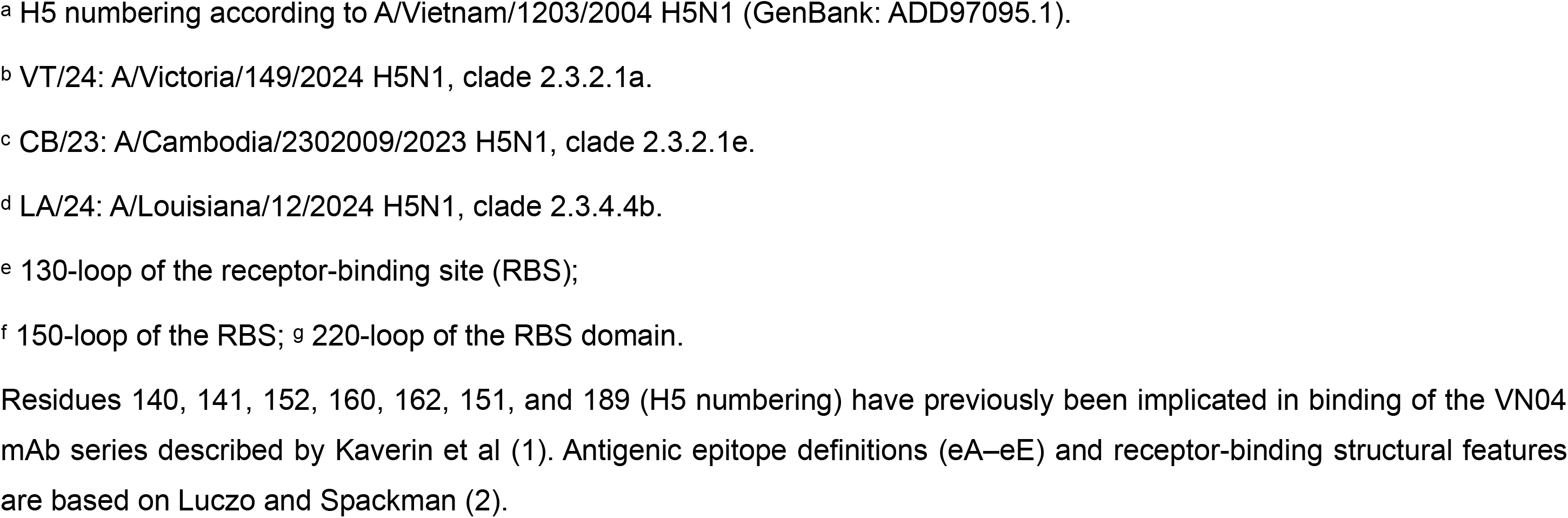
Amino acid substitutions within H5 HA antigenic epitopes of representative PR8-based H5N1 viruses relative to A/Vietnam/1203/2004 (VT04) H5N1.

**Supplementary Table S2.**
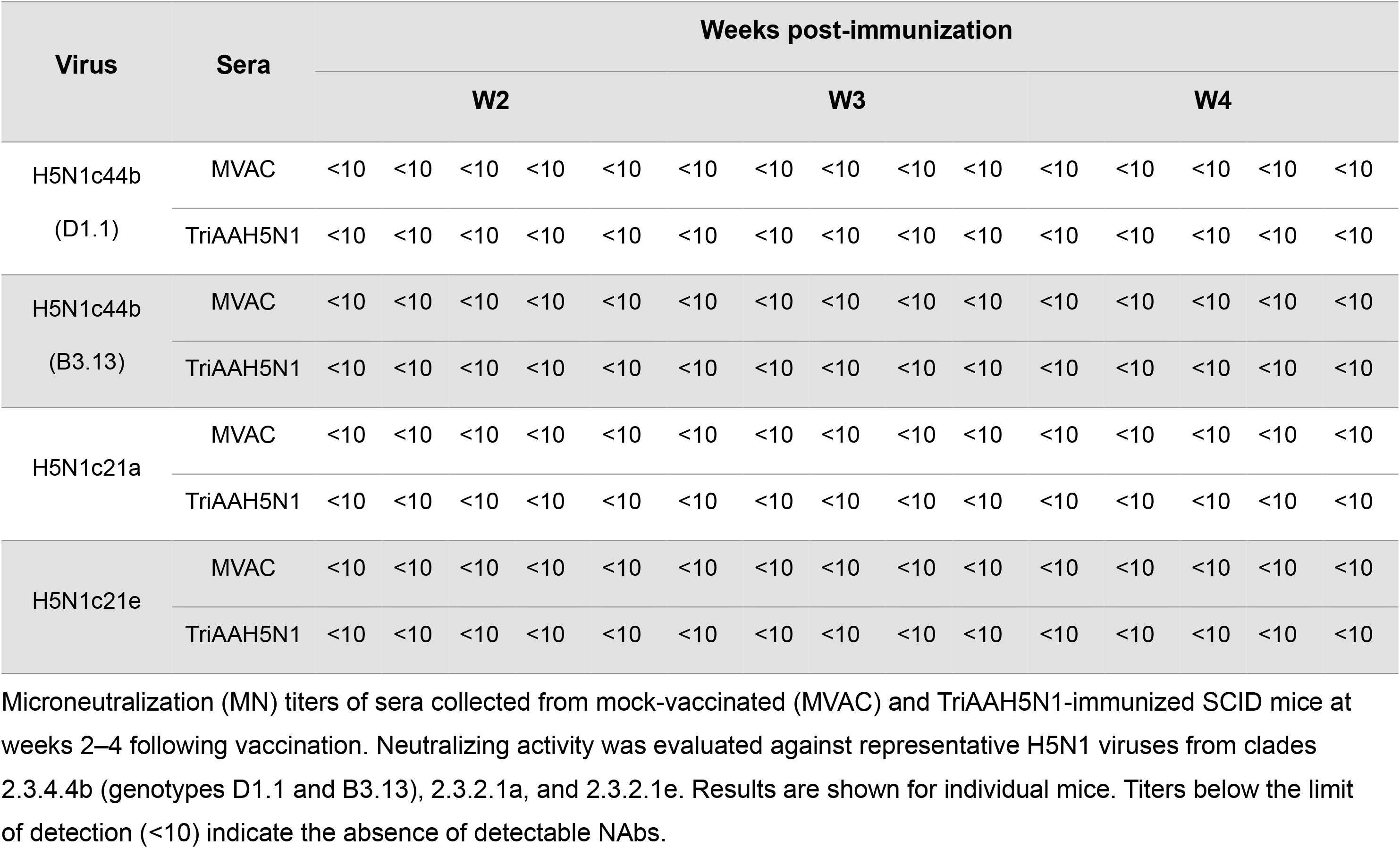
NAb responses in TriAAH5N1-immunized SCID mice following vaccination.

## References

1. Tyrrell CS, Allen JLY, Gkrania-Klotsas E. 2021. Influenza: epidemiology and hospital management. Medicine (Abingdon) 49:797–804.

2. Mostafa A, Abdelwhab EM, Mettenleiter TC, Pleschka S. 2018. Zoonotic Potential of Influenza A Viruses: A Comprehensive Overview. Viruses 10.

3. Mostafa A, Barre RS, Allué-Guardia A, Escobedo RA, Shivanna V, Rothan H, Castro EM, Ma Y, Cupic A, Jackson N, Bayoumi M, Torrelles JB, Ye C, García-Sastre A, Martinez-Sobrido L. 2025. Replication kinetics, pathogenicity and virus-induced cellular responses of cattle-origin influenza A(H5N1) isolates from Texas, United States. Emerging Microbes & Infections 14:2447614.

4. Mostafa A, Naguib MM, Nogales A, Barre RS, Stewart JP, Garcia-Sastre A, Martinez-Sobrido L. 2024. Avian influenza A (H5N1) virus in dairy cattle: origin, evolution, and cross-species transmission. mBio 15:e0254224.

5. CDC. 2026. A(H5) Bird Flu: Current Situation. https://www.cdc.gov/bird-flu/situation-summary/index.html. Accessed

6. Aguero M, Monne I, Sanchez A, Zecchin B, Fusaro A, Ruano MJ, Del Valle Arrojo M, Fernandez-Antonio R, Souto AM, Tordable P, Canas J, Bonfante F, Giussani E, Terregino C, Orejas JJ. 2023. Highly pathogenic avian influenza A(H5N1) virus infection in farmed minks, Spain, October 2022. Euro Surveill 28.

7. Domanska-Blicharz K, Swieton E, Swiatalska A, Monne I, Fusaro A, Tarasiuk K, Wyrostek K, Stys-Fijol N, Giza A, Pietruk M, Zecchin B, Pastori A, Adaszek L, Pomorska-Mol M, Tomczyk G, Terregino C, Winiarczyk S. 2023. Outbreak of highly pathogenic avian influenza A(H5N1) clade 2.3.4.4b virus in cats, Poland, June to July 2023. Euro Surveill 28.

8. Puryear W, Sawatzki K, Hill N, Foss A, Stone JJ, Doughty L, Walk D, Gilbert K, Murray M, Cox E, Patel P, Mertz Z, Ellis S, Taylor J, Fauquier D, Smith A, DiGiovanni RA, Jr., van de Guchte A, Gonzalez-Reiche AS, Khalil Z, van Bakel H, Torchetti MK, Lantz K, Lenoch JB, Runstadler J. 2023. Highly Pathogenic Avian Influenza A(H5N1) Virus Outbreak in New England Seals, United States. Emerg Infect Dis 29:786–791.

9. AbuBakar U, Amrani L, Kamarulzaman FA, Karsani SA, Hassandarvish P, Khairat JE. 2023. Avian Influenza Virus Tropism in Humans. Viruses 15.

10. WHO. 2026. Cumulative number of confirmed human cases for avian influenza A(H5N1) reported to WHO, 2003-2026, 22 January 2026. https://www.who.int/publications/m/item/cumulative-number-of-confirmed-human-cases-for-avian-influenza-a(h5n1)-reported-to-who--2003-2026--22-january-2026. Accessed

11. Nogales A, Villamayor L, Utrilla-Trigo S, Ortego J, Martinez-Sobrido L, DeDiego ML. 2021. Natural Selection of H5N1 Avian Influenza A Viruses with Increased PA-X and NS1 Shutoff Activity. Viruses 13.

12. Sutton TC. 2018. The Pandemic Threat of Emerging H5 and H7 Avian Influenza Viruses. Viruses 10.

13. Mostafa A, Nogales A, Martinez-Sobrido L. 2025. Highly pathogenic avian influenza H5N1 in the United States: recent incursions and spillover to cattle. npj Viruses 3:54.

14. Deng YM, Wille M, Dapat C, Xie R, Lay O, Peck H, Daley AJ, Dhanasakeran V, Barr IG. 2025. Influenza A(H5N1) Virus Clade 2.3.2.1a in Traveler Returning to Australia from India, 2024. Emerg Infect Dis 31:135–138.

15. Siegers Jurre Y, Xie R, Edwards Kimberly M, Byrne Alexander MP, Hu S, Wang R, Yann S, Sin S, Tok S, Chea K, Horm S, Rith C, Keo S, Pum L, Duong V, Auerswald H, Phou Y, Kol S, Spiegel A, Harvey R, Tum S, Sorn S, Seng B, Sengdoeurn Y, Chau D, Chin S, Hak M, Ieng V, Patel S, Thielen P, Claes Filip F, Lewis Nicola S, Ly S, Karlsson Erik A, Dhanasekaran V. 2025. Resurgence of Zoonotic Highly Pathogenic Avian Influenza A(H5N1) Virus in Cambodia. New England Journal of Medicine 393:1650–1652.

16. Chin S, Soputhy C, Seng H, Mom S, Sar B, Finlay A, Tan KR, Gould PL, Siegers JY, Karlsson EA, Olsen SJ, Uyeki TM, Davis WW, Chau D, Ly S. 2026. Investigation of a Family Cluster of Human Infections With Highly Pathogenic Avian Influenza A(H5N1) Virus, Clade 2.3.2.1e, in Cambodia, February 2023. Influenza Other Respir Viruses 20:e70231.

17. Barberis I, Myles P, Ault SK, Bragazzi NL, Martini M. 2016. History and evolution of influenza control through vaccination: from the first monovalent vaccine to universal vaccines. J Prev Med Hyg 57:E115–E120.

18. Krammer F. 2019. The human antibody response to influenza A virus infection and vaccination. Nat Rev Immunol 19:383–397.

19. Martinez-Sobrido L, Peersen O, Nogales A. 2018. Temperature Sensitive Mutations in Influenza A Viral Ribonucleoprotein Complex Responsible for the Attenuation of the Live Attenuated Influenza Vaccine. Viruses 10.

20. Nogales A, Martinez-Sobrido L. 2016. Reverse Genetics Approaches for the Development of Influenza Vaccines. Int J Mol Sci 18.

21. Sridhar S, Brokstad KA, Cox RJ. 2015. Influenza Vaccination Strategies: Comparing Inactivated and Live Attenuated Influenza Vaccines. Vaccines (Basel) 3:373–89.

22. Hoft DF, Lottenbach KR, Blazevic A, Turan A, Blevins TP, Pacatte TP, Yu Y, Mitchell MC, Hoft SG, Belshe RB. 2017. Comparisons of the Humoral and Cellular Immune Responses Induced by Live Attenuated Influenza Vaccine and Inactivated Influenza Vaccine in Adults. Clin Vaccine Immunol 24.

23. Blanco-Lobo P, Nogales A, Rodríguez L, Martínez-Sobrido L. 2019. Novel Approaches for The Development of Live Attenuated Influenza Vaccines. Viruses 11.

24. Nogales A, DeDiego ML, Martínez-Sobrido L. 2022. Live attenuated influenza A virus vaccines with modified NS1 proteins for veterinary use. Front Cell Infect Microbiol 12:954811.

25. Ilyushina NA, Haynes BC, Hoen AG, Khalenkov AM, Housman ML, Brown EP, Ackerman ME, Treanor JJ, Luke CJ, Subbarao K, Wright PF. 2015. Live attenuated and inactivated influenza vaccines in children. J Infect Dis 211:352–60.

26. Kitano T, Yoshida S. 2026. Effectiveness of live-attenuated and inactivated influenza vaccines against influenza in 2-17-y-old children, United States, 2022-2025. Hum Vaccin Immunother 22:2618341.

27. Wang L, Adolphus C, Wang J, Hossain J, Currier M, Davis CT, Dugan VG, Wentworth DE, Zhou B. 2026. Development of pre-pandemic influenza candidate vaccine viruses for use in vaccine manufacturing. NPJ Vaccines 11.

28. Rodriguez L, Blanco-Lobo P, Reilly EC, Maehigashi T, Nogales A, Smith A, Topham DJ, Dewhurst S, Kim B, Martínez-Sobrido L. 2019. Comparative Study of the Temperature Sensitive, Cold Adapted and Attenuated Mutations Present in the Master Donor Viruses of the Two Commercial Human Live Attenuated Influenza Vaccines. Viruses 11.

29. Cox NJ, Kitame F, Kendal AP, Maassab HF, Naeve C. 1988. Identification of sequence changes in the cold-adapted, live attenuated influenza vaccine strain, A/Ann Arbor/6/60 (H2N2). Virology 167:554–67.

30. Blais-Savoie J, Halajian E, Nirmalarajah K, Banete A, Corredor JC, Kotwa JD, Lee Y, Raj S, Sharif S, Mideo N, Mubareka S. 2026. Examining the Threat of H5N1 Highly Pathogenic Avian Influenza to Human Health. Chest 169:947–957.

31. Kaverin NV, Rudneva IA, Govorkova EA, Timofeeva TA, Shilov AA, Kochergin-Nikitsky KS, Krylov PS, Webster RG. 2007. Epitope mapping of the hemagglutinin molecule of a highly pathogenic H5N1 influenza virus by using monoclonal antibodies. J Virol 81:12911–7.

32. Wang L, Wang J, Hossain J, Cooper HC, Adolphus C, Currier M, Atteberry G, Feng C, Kirby MK, Di H, Barnes JR, Maines TR, Williams TL, Barr JR, Chen LM, Tumpey TM, Donis RO, Davis CT, Dugan VG, Wentworth DE, Zhou B. 2026. Development and Characterization of Candidate Vaccine Viruses against High Pathogenicity Avian Influenza A(H5) Viruses for Rapid Pandemic Response. J Infect Dis doi:10.1093/infdis/jiag132.

33. Sanz-Muñoz I, Sánchez-Martínez J, Rodríguez-Crespo C, Concha-Santos CS, Hernández M, Rojo-Rello S, Domínguez-Gil M, Mostafa A, Martinez-Sobrido L, Eiros JM, Nogales A. 2025. Are we serologically prepared against an avian influenza pandemic and could seasonal flu vaccines help us? mBio 16:e03721–24.

34. Sanz-Muñoz I, Ciria-Gil CJ, Martín-Toribio A, Toquero-Asensio M, Sánchez-Martínez J, Rodríguez-Crespo C, Hernandez M, Barragan-Martin I, Landeras-Bueno S, Eiros JM, Elsayed AM, Martinez-Sobrido L, Nogales A. 2026. Neuraminidase-Based Cross-Protective Immunity against H5N1 Influenza Viruses in Humans. bioRxiv doi:10.64898/2026.06.07.730769:2026.06.07.730769.

35. Liew KY, Aziz DB, Chan YT, Tan CW, Tambyah P, Tan Y-J. 2026. Neuraminidase-inhibiting antibodies boosted by H1N1pdm infection cross-react differently with H5N1 of clades 2.3.4.4b and 2.3.2.1a. Emerging Microbes & Infections 15:2662076.

36. Brock N, Pulit-Penaloza JA, Belser JA, Pappas C, Sun X, Kieran TJ, Zeng H, De La Cruz JA, Hatta Y, Di H, Davis CT, Tumpey TM, Maines TR. 2025. Avian Influenza A(H5N1) Isolated from Dairy Farm Worker, Michigan, USA. Emerging Infectious Diseases 31:1253–1256.

37. Chen GL, Lamirande EW, Jin H, Kemble G, Subbarao K. 2010. Safety, immunogencity, and efficacy of a cold-adapted A/Ann Arbor/6/60 (H2N2) vaccine in mice and ferrets. Virology 398:109–114.

38. Jin H, Subbarao K. 2015. Live attenuated influenza vaccine. Curr Top Microbiol Immunol 386:181–204.

39. Li H, Bradley KC, Long JS, Frise R, Ashcroft JW, Hartgroves LC, Shelton H, Makris S, Johansson C, Cao B, Barclay WS. 2018. Internal genes of a highly pathogenic H5N1 influenza virus determine high viral replication in myeloid cells and severe outcome of infection in mice. PLoS Pathog 14:e1006821.

40. Huang X, Yu D, Pan L, Wu X, Li J, Wang D, Liu L, Zhao C, Huang W. 2025. Increase in H5N1 vaccine antibodies confers cross-neutralization of highly pathogenic avian influenza H5N1. Nat Commun 16:5517.

41. Alosaimi B, Al-Rawi HZ, Alahmadi RM, Al-Shouli ST, Alzahrani J, Alsubki R, Awadalla ME. 2026. An epitope-based peptide vaccine targeting influenza a elicits robust immune responses and induces protection in a mouse model. Scientific Reports doi:10.1038/s41598-026-60696-3.

42. Yang S, Niu S, Guo Z, Yuan Y, Xue K, Liu S, Jin H. 2013. Cross-protective immunity against influenza A/H1N1 virus challenge in mice immunized with recombinant vaccine expressing HA gene of influenza A/H5N1 virus. Virol J 10:291.

43. Le Sage V, Werner BD, Merrbach GA, Petnuch SE, O’Connell AK, Simmons HC, McCarthy KR, Reed DS, Moncla LH, Bhavsar D, Krammer F, Crossland NA, McElroy AK, Duprex WP, Lakdawala SS. 2024. Pre-existing H1N1 immunity reduces severe disease with bovine H5N1 influenza virus. bioRxiv doi:10.1101/2024.10.23.619881.

44. Restori KH, Weaver V, Patel DR, Merrbach GA, Septer KM, Field CJ, Bernabe MJ, Kronthal EM, Minns A, Lindner SE, Lakdawala SS, Le Sage V, Sutton TC. Preexisting immunity to the 2009 pandemic H1N1 virus reduces susceptibility to H5N1 infection and disease in ferrets. Science Translational Medicine 17:eadw4856.

45. Kamei K. 2023. Live attenuated vaccines in patients receiving immunosuppressive agents. Pediatr Nephrol 38:3889–3900.

46. Tiozzo G, de Roo AM, Hofstra HS, Gurgel do Amaral GS, Vondeling GT, Postma MJ, Freriks RD. 2025. Safety of live attenuated vaccines in immunocompromised individuals and pregnant women: a systematic literature review. Expert Review of Vaccines 24:1033–1046.

47. Karron RA, Talaat K, Luke C, Callahan K, Thumar B, Dilorenzo S, McAuliffe J, Schappell E, Suguitan A, Mills K, Chen G, Lamirande E, Coelingh K, Jin H, Murphy BR, Kemble G, Subbarao K. 2009. Evaluation of two live attenuated cold-adapted H5N1 influenza virus vaccines in healthy adults. Vaccine 27:4953–60.

48. Subbarao K. 2021. Live Attenuated Cold-Adapted Influenza Vaccines. Cold Spring Harb Perspect Med 11.

49. Babu TM, Levine M, Fitzgerald T, Luke C, Sangster MY, Jin H, Topham D, Katz J, Treanor J, Subbarao K. 2014. Live attenuated H7N7 influenza vaccine primes for a vigorous antibody response to inactivated H7N7 influenza vaccine. Vaccine 32:6798–804.

50. Talaat KR, Luke CJ, Khurana S, Manischewitz J, King LR, McMahon BA, Karron RA, Lewis KD, Qin J, Follmann DA, Golding H, Neuzil KM, Subbarao K. 2014. A live attenuated influenza A(H5N1) vaccine induces long-term immunity in the absence of a primary antibody response. J Infect Dis 209:1860–9.

51. Mostafa A, Kanrai P, Petersen H, Ibrahim S, Rautenschlein S, Pleschka S. 2015. Efficient Generation of Recombinant Influenza A Viruses Employing a New Approach to Overcome the Genetic Instability of HA Segments. PLOS ONE 10:e0116917.

52. Barre RS, Escobedo RA, Castro EM, Gazi M, Castro JD, Cupic A, Bayoumi M, Jackson N, Ye C, Nogales A, Platt RN, Carrion R, Jr., Anderson TJC, García-Sastre A, Mostafa A, Martinez-Sobrido L. 2025. A human H5N1 influenza virus expressing bioluminescence for evaluating viral infection and identifying therapeutic interventions. iScience 28:113402.

53. Barre Ramya S, Mostafa A, Chiem K, Pearl Rebecca L, Platt Roy N, Cupic A, Anderson Timothy JC, Knaus Ulla G, Albrecht Randy A, García-Sastre A, Kobie James J, Nogales A, Martinez-Sobrido L. 2025. Bioluminescent reporter influenza A viruses to track viral infections. Microbiology Spectrum 13:e02150–25.

54. WHO. 2010. Serological diagnosis of influenza by microneutralization assay. https://www.who.int/publications/i/item/serological-diagnosis-of-influenza-by-microneutralization-assay.

55. Mostafa A, Ye C, Barre RS, Shivanna V, Meredith R, Platt RN, Escobedo RA, Bayoumi M, Castro EM, Jackson N, Cupic A, Nogales A, Anderson TJC, García-Sastre A, Martinez-Sobrido L. 2025. A live attenuated NS1-deficient vaccine candidate for cattle-origin influenza A (H5N1) clade 2.3.4.4.b viruses. npj Vaccines 10:151.

## Supplementary data references

1. Kaverin NV, Rudneva IA, Govorkova EA, Timofeeva TA, Shilov AA, Kochergin-Nikitsky KS, Krylov PS, Webster RG. 2007. Epitope mapping of the hemagglutinin molecule of a highly pathogenic H5N1 influenza virus by using monoclonal antibodies. J Virol 81:12911–7.

2. Luczo JM, Spackman E. 2024. Epitopes in the HA and NA of H5 and H7 avian influenza viruses that are important for antigenic drift. FEMS Microbiol Rev 48.

